# Heritable non-genetic phenotypes are enriched for stress responses as a form of bet hedging

**DOI:** 10.1101/2024.10.15.618459

**Authors:** Spencer Grissom, Zachary Dixon, Abhyudai Singh, Mark Blenner

## Abstract

During manufacturing batches, Chinese hamster ovary (CHO) cells encounter critical levels of environmental stressors such as ammonia, lactate, and osmolality accumulation that can significantly reduce cell health and productivity. It is therefore crucial that stress adaptation and resistance be factored into cell line development (CLD). In this study, we employee population-based transcriptomic and differential gene expression analysis on stress-induced CHO cells to identify biomarkers displaying both heritable and stress-responsive properties. Using this workflow, 199 genes displayed transcriptional variability characteristic of a bistable system that formed four network communities of co-fluctuating genes. These communities were enriched in genes related to the regulation of apoptotic processes and gene expression/metabolic pathways. Seven genes were identified as promising biomarkers for engineering a stress-resistant phenotype. Genetic engineering methods may be employed in the future to bias clonal populations for higher stress tolerance to manufacturing stress, therefore increasing cell health and productivity in at-scale bioreactors.

## 1. Introduction

Mammalian cells, in particular Chinese hamster ovary (CHO) cells, are favored for the production of mAbs due to favorable growth properties, rapid adaptability, and human-like post translational modifications (PTMs)^1,2^. As is true for many immortalized cell lines, CHO cells are also characterized by transcriptional plasticity that results in genetic drift, instability, and subclonal heterogeneity^3,4^. The observed plasticity is due in part to genomic instability in the form of karyotype rearrangements, copy number variations (CNVs), and small mutations (SMs)/structural variants (SVs) from faulty DNA repair machinery^4–7^. Another, and often overlooked, factor in CHO cell’s transcriptional heterogeneity is the rapidly changing epigenome, which is characterized by post-translational modifications to the DNA, RNA, or histone proteins along with non-coding regulatory RNA^8,9^. These chemical modifications to DNA and histones give rise to both rapid and transient changes as well as heritable and long-term memory states, and are often reactionary to environmental stimuli^10^. The relationship between epigenetic marks and effects on gene expression has been explored in CHO to demonstrate the formation of distinct chromatin states characterized by unique combinations of histone modifications and DNA methylation patterns in response to variable culture conditions^11–13^. Epigenetic regulation of gene expression has the advantage of speed and reversibility in contrast to genetic mutations. The lifetime of the resulting epigenetic states can last anywhere from weeks in the case of DNA methylation to days for histone acetylation, providing a range of time-scales for inheritance of epigenetic states^14,15^ The memory of these states are further reinforced by the crosstalk between epigenetic modifications where the writers, erasers, and readers either recruit each other to form complexes, such as the polycomb groups, or recognize certain marks to localize additional modifications, such as those containing methyl-CpG-binding domains^16–18^.

The result of these epigenetic modifications are distinct phenotypes characterized by multiple transient semi-stable levels of gene expression that are inherited from parent to daughter cell. These rare expression patterns have been identified in several organisms and the lifetime varies on whether the system is truly epigenetically maintained and forms a bistable state or is the consequence of transient coordinated fluctuations^19–21^. These heritable gene states described above characterize a phenomenon referred to as an epigenetic switch that permits the transition between various molecular states or expression levels^22^. Genes or gene networks with heritable epigenetic states promote phenotypic diversity and plasticity to permit rapid population-wide adaptation to environmental stress^23,24^. The expression of stress-resistance genes imposes a metabolic burden that reduces cell-growth and siphons resources to confer no additional advantage or fitness when a stressor is absent^25^. When an environmental stressor is encountered, the presence of a bistable system formed from epigenetic diversity allows for rapid adaptation within the population compared to the sluggish pace of genetic adaptation and increases the fitness for the stress-tolerant molecular state. Developing an understanding of the gene networks that exhibit this pattern of heritability can help clarify stress resistance mechanisms and suggest engineering targets to improve stress tolerance in CHO cell lines.

A simple and elegant experimental method known as MemorySeq was developed by Shaffer et. al. for the purpose of identifying these heritable gene expression states^26^. This technique is reminiscent of the classical Luria-Delbrück experiment to investigate whether bacteriophage resistance developed because of selective pressure or from random, but constant, fluctuations of a resistant phenotype during cell-division^27^. After plating a small number of *E. coli* cells in parallel cultures across multiple generations, they found the development of a resistant phenotype occurred spontaneously in the absence of selective pressure and was inherited by daughter cells. MemorySeq similarly explores intermediate cellular memory states by closely monitoring the transcriptomic profile of single cell derived clonal populations. This tool is powerful for identifying rare heritable phenotypes or gene expression states that would otherwise be indistinguishable from noise if using single-cell RNASeq (scRNAseq) without tools to trace shared lineages^28^.

For biopharmaceutical production, an optimal master cell line is characterized by a rapid and robust growth rate, high production and secretion capacity, desirable human-like post translational modifications, such as N-linked glycosylation, and long-duration phenotypic stability^29,30^. The cell line development (CLD) process is lengthy, labor intensive, and expensive. Thousands of single-cell clones are characterized for growth kinetics, productivity, product quality, and cell line stability.^31,32^ The conditions for screening occur at small-scale environments that do not mimic the stressful manufacturing conditions encountered in high-density cell cultures in large-scale bioreactors^31,33,34^. Manufacturing-scale conditions are characterized by the accumulation of toxic metabolic byproducts, shear during sparging and mixing, high osmolality during pH maintenance, and significant spatial and temporal heterogeneity^35–39^. Two common toxic byproducts often measured during a manufacturing cycle are ammonia and lactate. Towards the end of a production batch, these can achieve concentrations as high as 20 mM and have been shown to reduce specific cell growth rates, reduce productivity, and influence N-linked glycosylation patterns^35,40–42^. It is imperative to include the ability to adapt to manufacturing-scale stress in the design considerations for what constitutes a suitable master cell line. Many studies into stress response pathways only consider phenotypic changes and differentially expressed genes (DEG). While these studies highlight shifting gene expression patterns, they do not indicate which genes are responsible for stress adaptation and resistance^35,40,43,44^. Additionally, efforts to minimize the deleterious effects of stress have been solely focused on engineering controls, such as alternative feeding/agitation strategies, or long-term adaptation rather than targeted cell line engineering for a tolerant phenotype ^45–50^. This project seeks to address this gap in knowledge through the identification of heritable biomarkers that could be selected or engineered in a rational design approach to confer a stress-tolerant phenotype. Biomarkers are a gene expression pattern that confers a distinct phenotype and are often identified using transcriptomic, proteomic, and metabolomic profiling^51^. Using MemorySeq, over 190 genes demonstrating intermediate heritability were identified and their complex interactions were explored. When compared to (DEG) caused by CHO cell manufacturing stresses, we found these heritable gene networks are significantly enriched in stress response genes characterized by regulation of the cell cycle, apoptotic processes, and stimuli detection. Comparing DEG observed in stressful media to unstressed MemorySeq data enabled identification of genes involved in the initiation of stress resistance phenotypes.

## 2. Results

### 2.1 MemorySeq identifies heritable gene expression states in CHO cells

Following an adapted version of the MemorySeq workflow originally developed by Shaffer et. al., a panel of genes associated with rare phenotypes characterized by significant variability, and therefore heritability, were identified^26^. To simplify the model and assumptions, the bimodal system considered here includes an abnormal expression state, such as an uncharacteristically “high” or “low” expression level, (“On” state) and the most abundant expression state (“Off” state). A slow rate of transitioning to the “On” state paired with a high probability of inheriting the “On” state characterizes a heritable gene expression profile. Fluctuations between the “On” state and the “Off” state that are rapid and stochastic with no correlation between daughter and parent cell characterize non-heritable gene expression states. In other words, if the lifetime of the “On” expression state is significantly longer than the characteristic time for cell-division, the rare phenotype will persist across multiple generations and be enriched in the progeny (**Figure 1A**). Importantly, it would prove difficult to differentiate these two patterns as inheritance using scRNAseq, which requires tracking population dynamics and tracing common lineages between cells. The appearance of a rare phenotype alone is indistinguishable between natural noise or the shift to a stable heritable phenotype (Supplemental Figure 1).

**Figure 1.**
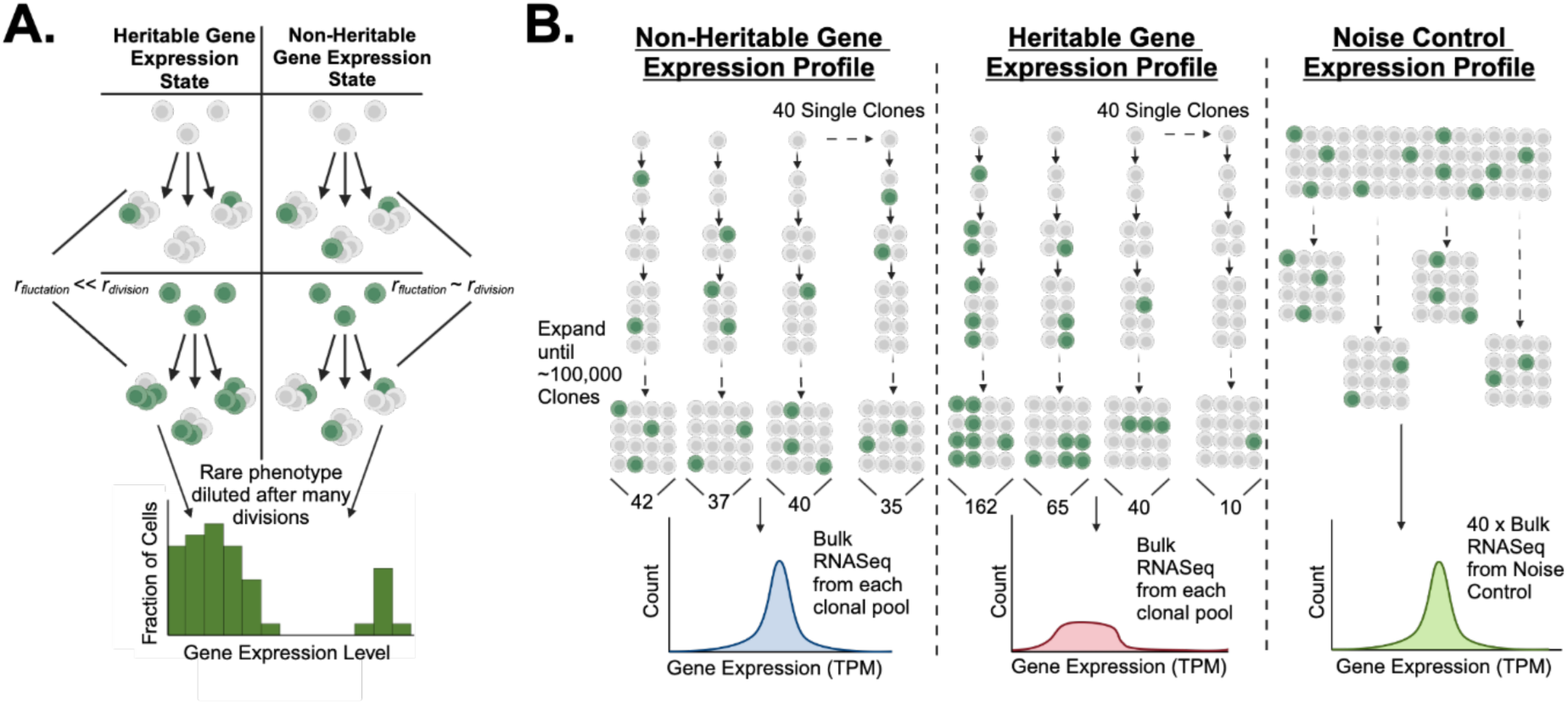
Summary of MemorySeq workflow and fluctuation analysis for identifying heritable gene states: Graphical principles for MemorySeq workflow (A.) Summary of heritable gene expression patterns for rare phenotypes. In this simplified model, green cells exhibit the rare-cell gene expression pattern and are surmised to be “On” while the white cells exhibit a bulk, lower average expression level and are said to be “Off”.(B.) MemorySeq workflow and basis for identifying heritable gene expression states. Starting from a bulk CHOZN^®^ GS^-/-^ Clone 23 passage flask, LDC techniques were used to seed sufficient wells to acquire n=40 single-cell clones. Comparison of bulk RNAseq samples between monoclonal derived cell pools and those from a bulk population forms the basis for how heritable genes are identified as heritable genes form a non-Gaussian expression distribution while non-heritable genes do not. Created in BioRender.com

In MemorySeq, around 40 single-cell populations of the CHOZN^®^ GS^-/-^ Clone 23 strain were seeded using limited dilution cloning (LDC) techniques to form the MemorySeq clones. These monoclonal populations were expanded until there were roughly 100,000 cells, reflecting 16-17 generations of growth or over 3-4 weeks. At this time, the whole RNA content of the MemorySeq clones was extracted and sequenced alongside 40 RNA samples collected from 100,000 bulk cells in a standard passage flask to constitute the noise control, where there is no shared lineage in the resulting population. For protein-coding genes, there are two expected distributions of gene expression. The first is a roughly normal distribution centered near the noise control average and reflects non-heritable expression states as the rapid fluctuations maintain a roughly constant number of high expressing clones. This distribution displays relatively low variability in expression. The other is typically a positive or negative-skew distribution characterized by high variance with an elongated tail. The variance and skew are explained by the lineage-dependent expression level as the generation at which a parent cell first transitions into the rare, “On” expression state influences the final expression level of the progeny. If the transition occurs early and the “On” state correlates with higher than average expression, then a higher proportion of the progeny will remain in a high expression state towards the end of growth compared to a MemorySeq clone that transitioned late or never transitioned at all. The resulting distribution would appear right skewed (**Figure 1B**.). Noise control samples are expected to be tightly distributed as there is no common lineage between cells in the sample and bulk expression is biased towards the average expression level.

### 2.2 MemorySeq identifies 199 heritable gene states in CHO cells

After all RNA samples were collected and submitted for sequencing, there were 38 MemorySeq samples and 40 noise control samples left after quality control. Of the 32,428 initial gene counts measured, only protein-coding genes with a minimum expression level of 2.5 transcripts per million (TPM) were included in the analysis as low expressing genes display greater variability that may skew downstream analysis^26^. Metrics of variation were calculated for the remaining 10,106 genes. Heritability is marked by significantly greater variability and skew compared to rapidly fluctuating non-heritable gene states (**Supplemental Figure 2**). A Poisson regression model accurately fits the relationship between coefficient of variation in TPM and the log_2_(Mean_TPM_) for most non-heritable genes. Genes that deviate significantly from the fit are those that have a uniquely high variance for its corresponding expression level. In this work, heritable gene expressions states were identified as those genes in the 98^th^ percentile for residuals compared to the Poisson regression model (**Figure 2A**)^26^. A gene displaying no tendency to inherit a rare gene expression state would possess a roughly 1:1 ratio of the CV between the MemorySeq samples and the noise control. The overall variance observed in MemorySeq clones is innately higher than a noise control sample due to the bias of cell-to-cell variability, yielding a ratio of 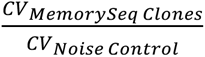 greater than unity. Despite this confounding noise, the Poisson regression method for quantifying heritability successfully identifies genes with disproportionally higher variance compared to this noise control counterpart (**Figure 2B**). There were 15 genes within the noise control that exceeded the residual threshold and any redundancy within the MemorySeq clones were removed as the natural variability of these genes confounds variability due to heritable principles. Residuals were compared to the critical threshold and 199 heritable gene states were identified. These genes appeared to be spread out in clusters across all 21 chromosomes with some common regulation patterns (**Supplemental Figure 2**) and displayed the characteristic “smear” or positive skew distribution in MemorySeq clones expected for heritable genes compared to the narrow and low variation noise control (**Figure 2C**). MemorySeq clones with higher expression levels for these heritable gene states are hypothesized to have transitioned into the rare “high” expression state earlier than the average clone.

**Figure 2.**
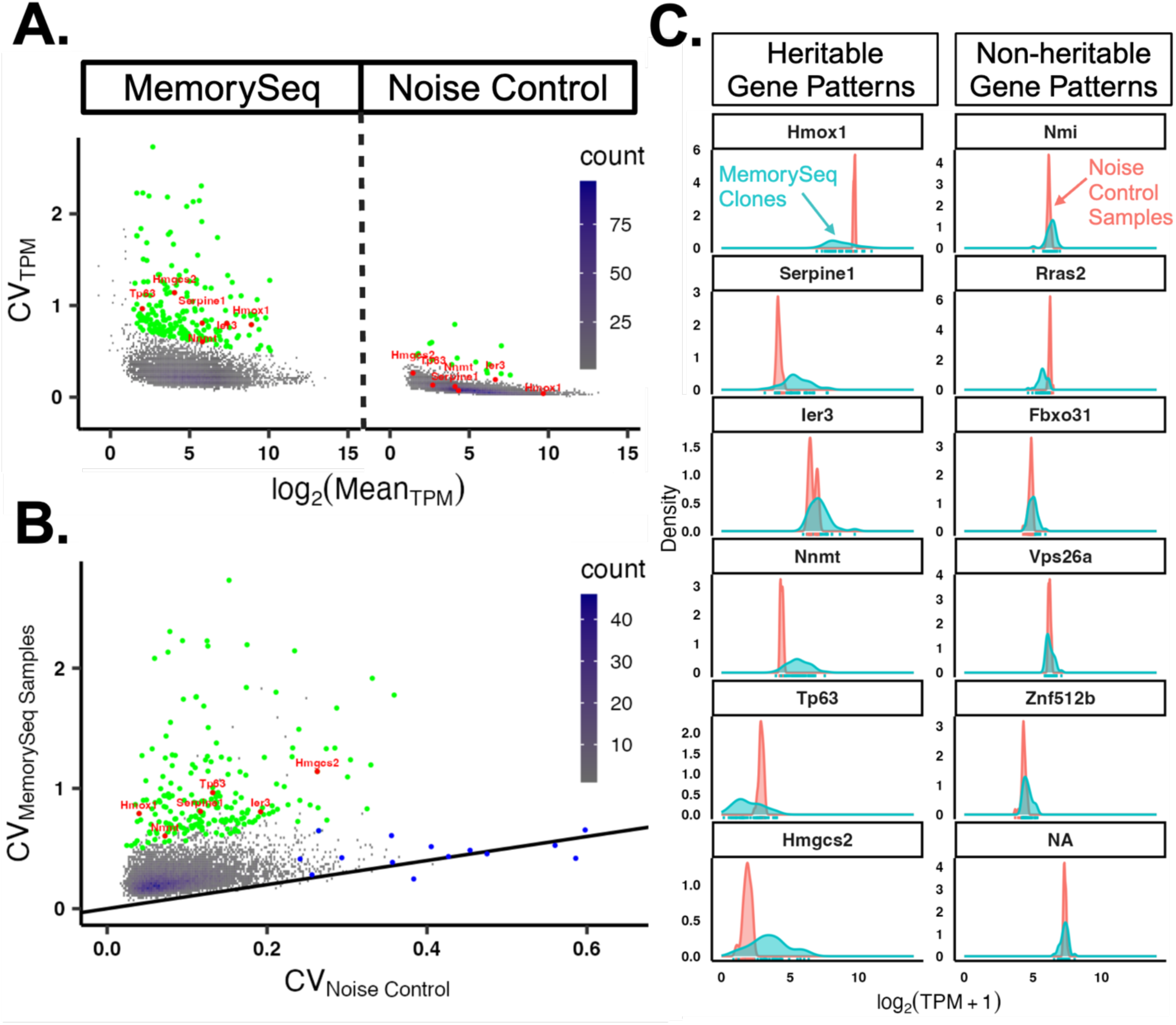
Application of MemorySeq identifies 199 heritable gene states in CHO cells: Visualization of heritable and non-heritable gene expression states (A.) Coefficient of variation for TPM versus log normalized average TPM for noise control and MemorySeq samples. Each dot represents one of 10,106 different genes meeting certain requirements or a cluster of genes. Upon fitting a Poisson regression distribution to each group, the genes that were in the top 2% of residuals and had an average log_2_(TPM) of 2 or greater are highlighted as green and considered to have a heritable gene expression profile. Any overlapping genes that were in the top 2% residuals and in the noise control were removed from consideration due to inherit biological noise. Genes of interest, to be discussed later, are marked in red. (B.) Coefficient of variation for TPM for MemorySeq Samples versus noise control. Each dot represents a different gene or cluster of genes. Green dots are those in the top 2% of residuals from the Poisson regression fit from the MemorySeq Samples while the blue dots are those in the top 2% of residuals from the noise control. (C.) Sample histograms for gene expression for 6 heritable (Left) and 6 non-heritable (Right) gene expression patterns. The blue curve corresponds to the distribution of MemorySeq clone expression and the pink is the distribution of noise control expression.

### 2.3 smRNA-FISH verifies heritable properties of *Hmox1* and *Ier3* by correlating relatedness to proximity

To verify the heritability principles for some of the physiologically relevant genes identified as heritable, small molecule RNA fluorescence *in-situ* hybridization (smRNA-FISH) was completed for *Hmox1* and *Ier3*. These genes were chosen as they had the highest basal expression, making signal intensity easier to capture, from the seven genes of interest identified as physiologically relevant further in this work. To measure the shared lineage between cells across multiple cell-divisions, cells were first attached to a fibronectin-coated coverglass to promote cellular adhesion and grown to 60-70% confluence. In doing so, the proximity of cells with respect to one another is good approximation for their relatedness or common lineage. As cells begin to transition to the high expression state, they serve as a nucleus for a larger, contiguous population of highly expressing cells if they are heritable (**Supplemental Figure 3**). This pattern of outgrowth can be captured visually by hybridization to fluorescently labeled RNA probes corresponding to some of the heritable genes of interest. Highly expressing cells were rare in the bulk population, but several pockets of such cells were observed for *Hmox1* and *Ier3*. Similar methods employed by Shaffer, et. al. utilized single-cell spatial analysis to verify that these patches of high expressing clones are unlikely to occur through random distribution, but originate from a shared lineage given their spatial density^1^.

### 2.4 Network community identification of heritable gene states reveals four co-fluctuating communities with shared biological processes

Complex phenotypes are rarely the result of a single gene, but rather the simultaneous co-fluctuation of network communities. To further elucidate the relationships between the 199 heritable gene expressions states, a Pearson correlation coefficient was calculated between all pairwise comparisons (**Figure 3.A.**). The plurality of genes exhibited a strongly positive correlation, indicating a co-regulation that activates gene expression. To investigate how sensitive these correlations were to outliers, the Cook’s distance of strongly correlated pairs was measured to determine if any biological replicate consistently skewed the correlations. Some outliers are implicitly expected in the data as an early transition to the rare “high” expression state yields a heavily skewed distribution. However, if an outlier is considerably larger or smaller than the average, it suppresses the variability captured in the remaining MemorySeq clones and dominates the magnitude of the variance and correlation. This can shift a gene that otherwise appears non-heritable to a heritable classification. The MemorySeq-2 clonal lineage was identified as a consistent outlier in Cook’s distance analysis and was removed from the data set to yield 37 MemorySeq replicates and 40 noise control replicates (**Supplemental Figure 4**).

**Figure 3.**
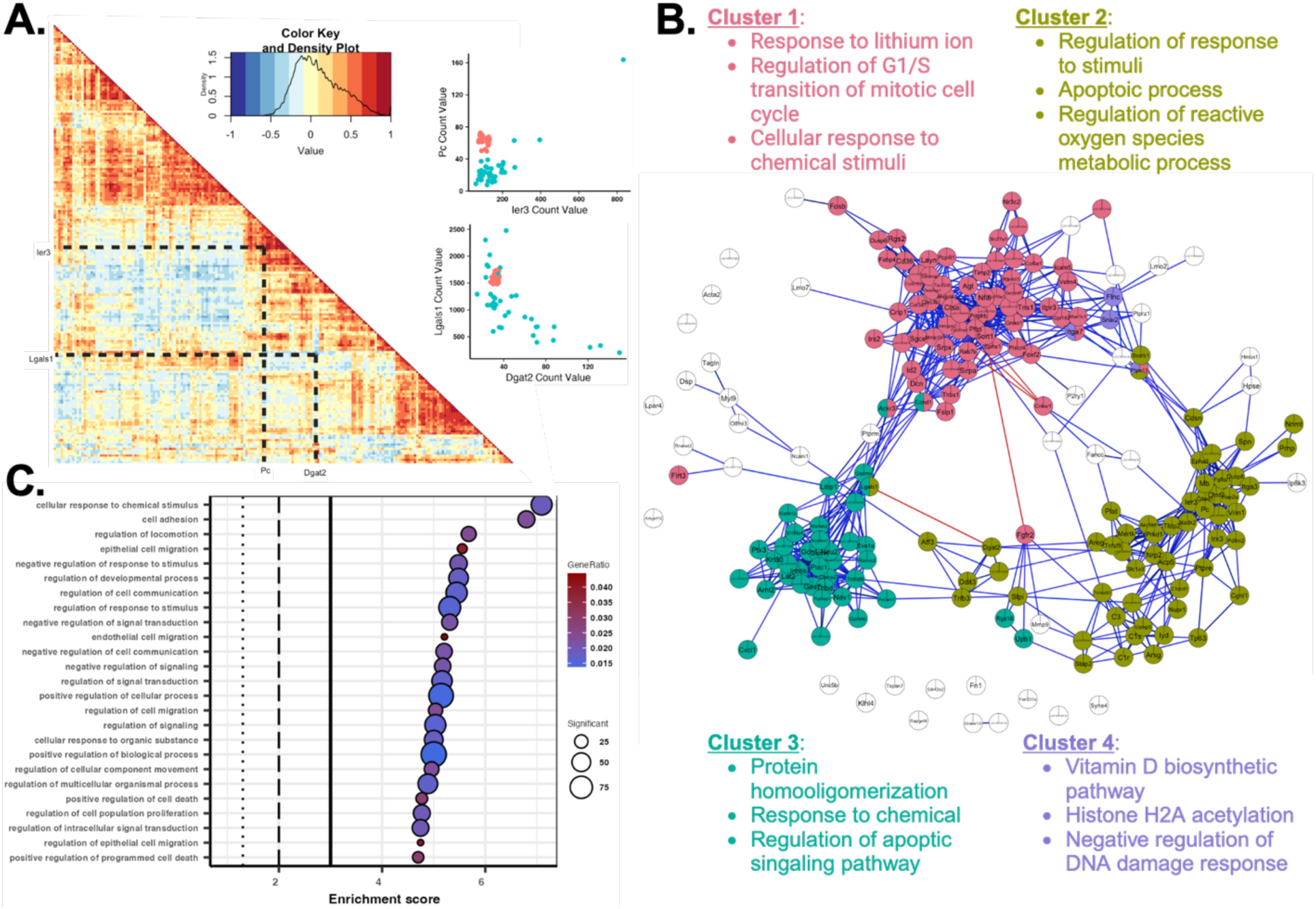
Network community identification of heritable gene states reveals four co-fluctuating communities with shared biological processes: (A.) Pearson pairwise correlation matrix for all 199 heritable gene states to highlight expression co-fluctuations between gene pairs. Red or positive correlations signify the mutual increase in expression levels between a gene pair (Example between Pc and Cited1 shown in bottom right). Blue or negative correlations signify the reduced expression of one gene coinciding with the enhanced expression of another gene (Example between Lgals1 and Dgat2 shown in top right). (B.) Product of k-clique percolation for network community identification of the 199 heritable gene states for k=4 and I=0.68. 4 distinct communities identified, differentiated by different colors along with GO terms enriched within the individual communities. Unlinked genes are shown in white and did not meet the k-1 requirement for inclusion within other networks. (C.) Dot plot visualizing the 25 most enriched GO terms across all 199 heritable gene states. Size and color intensity reflect the number of genes contained with each GO term and the corresponding gene ratio. Dotted lines represent a p-value of 0.05, 0.01, and 0.001 from left to right.

Network community identification from the correlations of pairwise co-fluctuating genes was carried out through a *k*-clique percolation method (*k*-CPM). The output of *k*-CPM is the identification of overlapping network communities of strongly co-fluctuating gene expression states that may outline a unique phenotype^52,53^. This method simplifies the network by assigning each gene as an independent node connected to others by the pairwise correlation. The size of the communities is defined by the value of *k* and the minimum correlation that connects nodes is defined by the threshold, *I*. A value of *k* = 4 was chosen to balance the number of internally connected communities and the cohesion of the community^26,54,55^ and the optimized *I* was identified to be 0.68 (**Supplemental Figure 5**). There were three large communities and one small community identified under these conditions (**Figure 3B**.). GO enrichment analysis was employed to identify and assess overrepresented biological processes unique to each network community. Of interest are processes related to cell health and fitness during stress, which are closely related to cell cycle/differentiation/apoptosis, metabolism, transcriptional/translational control, and response to internal and external changes^37,56,57^. Enriched across the entire heritable gene pool were 11 children GO terms relating to the response to stimuli, including the response to chemical, external stimuli, and stress. Likewise, 5 children terms relating to apoptotic processes were enriched, most notably the regulation of apoptosis. Other noteworthy mentions include regulation of cell adhesion, cell communication, and cellular compartmentalization (**Figure 3.C.**). These two biological processes constitute the immediate detection of external stress and the resulting response for unresolved stress when cytoprotective forces are insufficient to promote cell survival^58^. In addition to genes relating to these two biological processes and specific to certain community networks identifier, there were other noteworthy enriched terms including cell cycle/proliferation, regulation of gene expression due to epigenetics/protein activity, and cell differentiation (**Supplemental Figure 6**). These networks may outline orthogonal mechanisms for how CHO develops a stress-resistant phenotype. The rare and random fluctuation of a stress-resistant gene that displays heritable characteristics into a high expression state would poise a small population of cells to be ready to combat the deleterious effects of the stressors while conferring favorable growth kinetics or survivability that are inherited by daughter cells. In this specific context, the mechanism behind the co-fluctuations is not well understood, but may be mutually linked to the expression or activation of common upstream regulatory factors, or related to the spatial proximity of the genes as measured through Hi-C sequencing methods^59–61^. Provided the co-fluctuation can be replicated, then the transition of any one of the genes within the network community may cascade until a comprehensive stress-resistant phenotype develops.

### 2.5 Elevated ammonia, lactate, and osmolality influences CHO growth rate and specific productivity during fed-batch

While the MemorySeq data indicates genes linked to a stress-resistance phenotype, it does not specify which genes correspond with resistance to which stressors. To explore this relationship, CHOZN^®^ GS^-/-^ Clone 23 was grown in production fed-batch flasks in the presence of biomanufacturing-relevant levels of ammonia, lactate, and osmolality following a 2^3^ factorial design of experiment (DOE) given these chemicals hinder cell growth kinetics and reduces overall cell health^35,41,62^ (**Figures 4A and 4B**). These stressors were chosen as they could be chemically manipulated in a fed-batch shake flask environment unlike shear or gas sparging. Ammonia accumulates as a waste product from cellular metabolism, primarily glutamine metabolism. At concentrations as low as 2-3 mM, deleterious effects such as inhibited cell growth or reduced integral viable cell density (IVCD), reduced cell specific productivity (qP), and worsened product quality attributes begin^35,63^. In the fed-batch DOE, ammonia was determined to have significantly reduced IVCD, but effects to qP were not as powerful (analysis of variation or ANOVA test, *p* < 0.05). Under fed-batch conditions, ammonia concentrations reach as high as 10-20 mM^35,40,42,64^. Compared to the control flask, ammonia concentrations of 10 mM resulted in a 50 ± 19% reduction in IVCD without reducing qP. (**Figure 4C)**. Unlike ammonia, lactate often has both an accumulation phase during glucose metabolism and a consumption phase. Consumption typically begins during periods of low glucose and high lactate concentrations seen during stationary phase growth. High concentration of lactate is toxic to cells and reduces productivity, while lower concentrations can support cell growth as an alternative carbon source. The toxicity of lactate is often concomitant with reduced pH and osmolarity effects. Prior to lactate consumption phases in bioreactors, lactate often hits concentrations as high as 15-20 mM^41,44,46,65,66^. At concentrations of 15 mM, there was no deleterious effect on cell growth observed, but qP was increased 32 ± 14%. At this concentration, the lactate may not have been toxic and instead provided an alternative substrate for growth and production (**Figure 4C**). Overall, there was no statistically significant effect on qP, but a strong positive effect on IVCD was observed (ANOVA test, *p* < 0.05). Finally, osmolality increases because of the secretion of ionic waste products, during pH maintenance through alkaline salts, and during continuous feed of nutrients during fed-batch conditions. Hyperosmolar conditions often results in enhanced productivity at the cost of cell volume increase, cell-cycle arrest, and reduced proliferation rates^62,67–69^. Consistent with literature, hyperosmolar conditions reduced IVCD by nearly 41 ±14% while improving qP by 65 ± 16%, both of which were statistically significant (ANOVA test, *p* < 0.05) (**Figure 4D**). Interactions between the different stressors highlight a few trends. Ammonia’s deleterious effect on growth was mitigated by the presence of lactate, but it did consistently reduce qP. This perhaps suggests a shift from producing to stabilizing growth and cell health.

**Figure 4.**
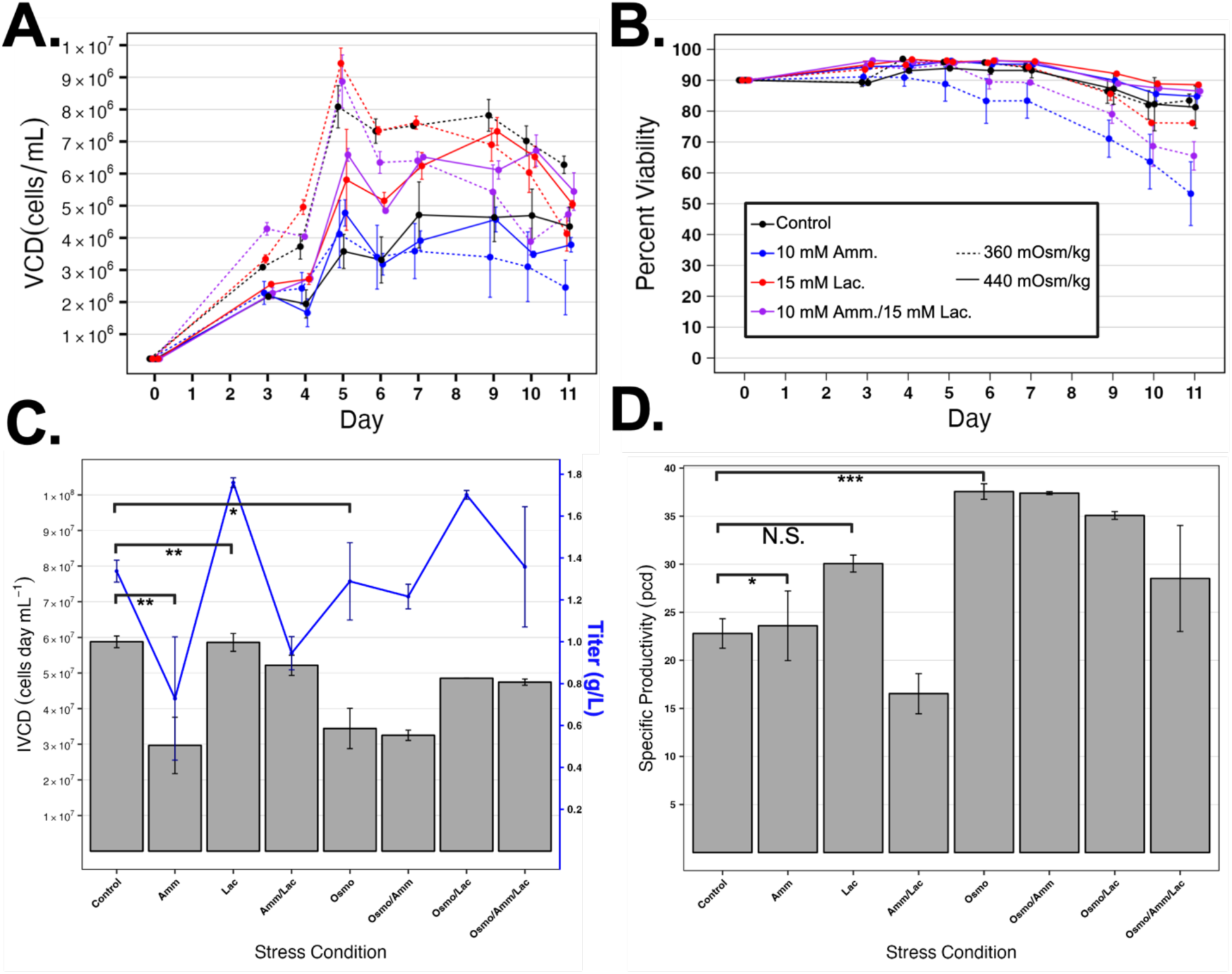
Elevated ammonia, lactate, and osmolality influences CHO growth rate and specific productivity during fed-batch: Manufacturing stress fed-batch culture performance. Ammonia, lactate, and osmolality stress conducted at 10 mM, 15 mM, and 440 mOsm/kg respectively (base osmolality was 360 mOsm/kg). All conditions tested at n=2 and error bars represent standard error of mean (SEM). (A.) Percent viability data and (B.) viable cell density tracked daily for fed-batch flasks. (C.) Overall cell growth and production for n=2. Gray bars represent integrated viable cell density (IVCD) and blue dots represents the mAb titer from the final day of the fed-batch culture. Error bars reflect the SEM. Significance lines represent the results of the pairwise ANOVA results where *, **, and *** represent p-values of 0.05, 0.01,and 0,0001 respectively. (D.) Cell specific productivity in units of picograms per cell per day. Error bars reflect SEM.

### 2.6 Combination of MemorySeq fluctuation analysis and differential gene expression analysis highlights promising biomarkers for cell-line engineering

The intersection between differentially expressed genes (DEG) and genes identified as heritable may suggest a possible route for which the stress-resistant phenotype developed. Day 5 RNA samples were collected from the high ammonia, lactate, osmolality, and combination of the three for differential gene expression analysis (DGEA). While all samples displayed a shift in gene expression, in agreement with the growth, productivity, and viability data, there was a muted difference between high lactate and the control (**Supplemental Figure 7**). The combinatorial stress condition had the most DEGs at 1275 and high lactate had the least at 155 DEGs. For each stress condition, the ratio of DEGs that were heritable to total DEGs significantly exceeded the expected ratio (Chi-squared goodness of fit test, *p*<0.001) given 10,106 genes from DGEA met filter parameters and 199 of these genes were considered heritable (**Table 1**). This finding reinforces that the heritable gene expression states significantly overlap with stress responsive genes. GO enrichment analysis was conducted on the overlapping DEG and heritable genes for each stress condition. Between all four conditions, biological processes related to response to stimuli, metabolism, and regulation of apoptosis/cell-cycle were over-represented. These biological processes outline the recognition of an environmental stress, shift in intracellular conditions, and the phenotypic outcome of cell health (**Supplemental Figure 7**).

**Table 1:** Relative enrichment of genes with heritable expression states within the DEG pool. Provided there were 199 heritable gene states out of the 10,106 genes considered, the expected ratio would be roughly 2%. Significant difference observed between the observed ratio and expected (Chi-Squared Goodness of Fit Test, p<0.001)

| **Stress**<br>**Condition** | **Observed**<br>**<u>*DEG and Heritable*</u>**<br>***DEG*** | **Expected**<br>**<u>*DEG and Heritable*</u>**<br>***DEG*** |
| --- | --- | --- |
| Ammonia | $\frac{61}{704}$ | $\frac{14}{704}$ |
| Lactate | $\frac{17}{155}$ | $\frac{3}{155}$ |
| Osmolality | $\frac{77}{1147}$ | $\frac{23}{1147}$ |
| Combination | $\frac{96}{1275}$ | $\frac{25}{1275}$ |

Genes with biological processes related to stress resistance and cell-health were extracted for all heritable genes. Inspection of these vital biological processes revealed that 30-50% of heritable genes related to the downregulation of apoptosis were significantly overexpressed in the stressed cultures except for lactate. Genes related to the upregulation of apoptosis were mixed, signifying genes that relate to the mechanism for resistance as well as the source of worsened cell health. Of the four heritable genes related to the endoplasmic reticulum unfolded protein response (ER UPR), *Atf3*, *Ccnd1*, and *Ddit3* were downregulated across the four stress conditions (**Figure 5**)^70,71^. Each of these genes play a role in the UPR and other stress response pathways to combat ER or external stress signals through induction of apoptosis^72–75^. Their downregulation suggests another mechanism for how resistance develops in biopharmaceutical producing CHO cells where external stress compounds upon the metabolic burden of mAb production.

**Figure 5:**
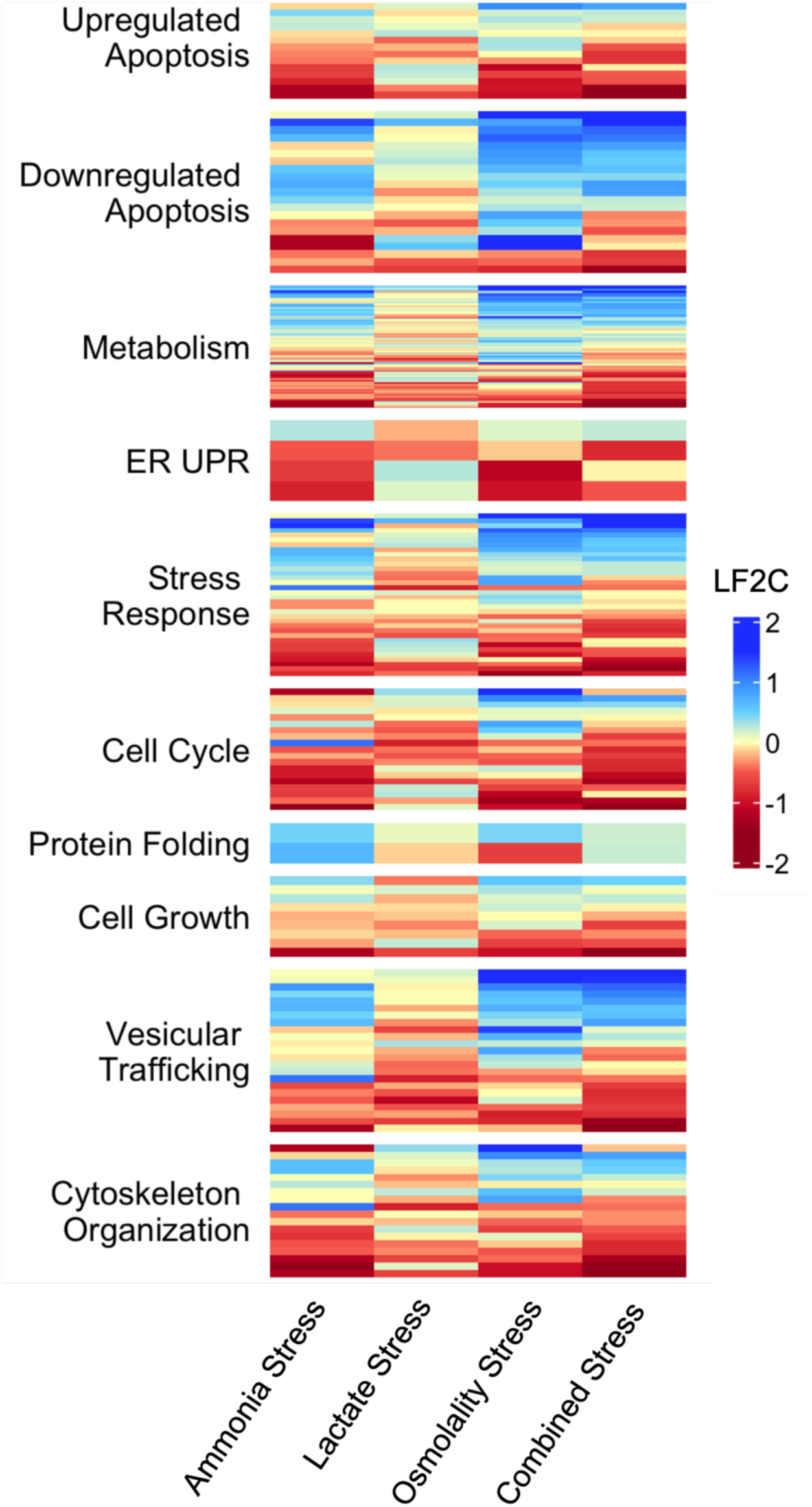
Combination of MemorySeq fluctuation analysis and differential gene expression analysis highlights promising biomarkers for cell-line engineering: Relative gene expression of genes with heritable expression states for common stress response pathways. Heatmap of log fold_2_ change (LF2C) of heritable gene expression states compared to control fed-batch flask for high ammonia, lactate, osmolality, and combination.

While not a necessity for a suitable biomarker, the lifetime of these heritable expression states, assuming a 1% frequency of the rare phenotype in bulk populations, was estimated to be between 5 and 10 generations for all genes identified in this work (See **Supplemental File 3**). The lifetime, which monotonically increases with measured coefficient of variance, provides some insight on gene states that may have more robust mechanisms for maintaining rare phenotypes and are perhaps more key to the early emergence of resistance pathways. From this analysis, 7 promising biomarkers for resistance to manufacturing stressors were identified (**Table 2**). This list includes genes that were upregulated across many of the stress conditions, were identified as heritable, and have been previously characterized as influencing apoptotic processes. Some of which, such as *Hmox1* and *Nnmt*, have been directly shown to display cytoprotective properties in response to oxidative and other sources of stress. Other genes, such as *Ier3*, *Tp63*, and *Atf3*, more ambiguously influence apoptosis and appear to be cell line dependent.

**Table 2:** Overview of promising biomarkers that exhibit heritable properties and play a role in stress adaptation or response. Summarizes LF2C for different stress conditions and the relevant biological function (* indicates p>0.05 and |L2FC|<0.58, ** indicates p<0.05 and |L2FC|<0.58)

| Gene Symbol | Stress Condition | Log<sub>2</sub>(FC) | Generation Lifetime of ON State | Relevant Biological Processes | References |
| --- | --- | --- | --- | --- | --- |
| *Hmox1* | Ammonia | 1.38 | 5.50 | Provides cytoprotective forces against cellular and oxidative stress. Active in cell proliferation and anti-apoptotic processes | 103–105 |
|  | Lactate | 0.68 |  |  |  |
|  | Osmolality | 0.83 |  |  |  |
|  | Everything | 1.64 |  |  |  |
| *Serpine1* | Ammonia | 0.63 | 5.53 | Anti-apoptotic processes and enhanced proliferation upon activation of Akt and ERK signaling pathway. Also shown to correlate with higher productivity in CHO. | 106–110 |
|  | Lactate | -0.33** |  |  |  |
|  | Osmolality | 0.29** |  |  |  |
|  | Everything | 0.84 |  |  |  |
| *Ier3* | Ammonia | 0.03* | 5.48 | Early response gene that responds to environmental stimuli and is regulated by ERK signaling pathway. Regulates apoptosis in a cell-dependent manner | 111–114 |
|  | Lactate | 0.15* |  |  |  |
|  | Osmolality | 2.17 |  |  |  |
|  | Everything | 1.78 |  |  |  |
| *Nnmt* | Ammonia | -1.15 | 4.96 | Reduce apoptosis through the mitochondria-mediated pathway. Silencing inhibits cell proliferation. Provides resistance to oxidative stress | 115,116 |
|  | Lactate | 0.19** |  |  |  |
|  | Osmolality | -0.17** |  |  |  |
|  | Everything | -0.80 |  |  |  |
| *Tp63* | Ammonia | -1.33 | 5.95 | Acts as a sequence specific DNA binding transcriptional activator or repressor. May be required in conjunction with TP73/p73 for initiation of p53/TP53 dependent apoptosis. Cell-line dependent behavior | 117,117,118 |
|  | Lactate | -0.24* |  |  |  |
|  | Osmolality | -0.66 |  |  |  |
|  | Everything | -2.07 |  |  |  |
| *Hmgcs2* | Ammonia | 0.01* | 6.35 | Catalyzes step in ketogenesis, condensing acetyl-CoA to acetoacetyl-CoA to form HMG-CoA. Plays a role in cholesterol biosynthetic pathway and expands secretory framework | 119,120 |
|  | Lactate | -0.70 |  |  |  |
|  | Osmolality | 0.64 |  |  |  |
|  | Everything | -0.92 |  |  |  |
| *ATF3* | Ammonia | -0.90 | 5.03 | Functions as an oncogene in prostate cancer by enhancing cell proliferation, gene regulation by recruiting p300, and stress response. Appears to co-localize with p53 | 71,72,121 |
|  | Lactate | 0.13* |  |  |  |
|  | Osmolality | -0.99 |  |  |  |
|  | Everything | -0.54** |  |  |  |

## 3. Discussion

Biopharmaceutical manufacturing stress is an often-unavoidable obstacle for CLD and production at large-scale, and can significantly alter cell line performance with regards to cell health, productivity, and product quality. This work sought to develop a more comprehensive understanding of how random intermediate memory states may give rise stress-resistant phenotypes and identify biomarkers that improve cell performance under such conditions. For years, large-scale transcriptomic analyses have been utilized to characterize population-based gene expression and monitor, to a lesser extent, cell-to-cell heterogeneity. However, the two most popular transcriptomic profiling tools, bulk RNAseq and scRNAseq, fail to capture the generational variability and the development of novel phenotypic patterns^76,77^. Bulk RNAseq is well-suited for measuring broad changes in gene expression, but due to an intrinsic sampling bias when collecting from a large population, it can under or over predict rare phenotypes ^77,78^. By contrast, scRNAseq samples the transcriptomic profile from individual cells isolated from the bulk population. The single-cell resolution of gene expression provides significant information on cellular heterogeneity but is limited on sample size, is difficult to connect to multi-omic datasets, and cannot track the shared lineage between cells without the use of extensive bar-coding^28^. Efforts to characterize single-cell heterogeneity in CHO have been mired by the intrinsic noise contained within. One transcriptomic study using scRNAseq identified an increase in heterogeneity as a result of increased population doubling levels (PDLs), but were only able to identify one gene, enolase, out of thousands as the sole biomarker that characterized distinct subpopulations^79^. Likewise, another scRNAseq study considering CHO-K1 and HEK293 cells and their storage conditions identified a set of highly variable genes, using a similar heuristic of CV scaled to transcriptional level used in this work, to characterize the heterogeneity of clones and identify any hidden subpopulations^80^. While they were unable to find any marker genes for subpopulations, the GO terms of the highly variable genes they did collect shared similar themes as the ones found in this work for heritable genes. This includes response to stimuli, cell-cycle regulation, and cell differentiation^80^. The true shortcoming of these methods is the inability to connect the shared lineage of the cells in either bulk or scRNAseq methods. By expanding cells in parallel, as described in the MemorySeq workflow, each of the monoclonal derived cell pools have a shared internal lineage. The heterogeneity in gene expression that is measured is therefore a reflection of intermediate transcriptional fluctuations that are time dependent. If the monoclonal expansion were carried out well beyond 100,000 clones, the RNAseq profile would gradually shift towards the bulk RNAseq result captured from the noise control.

Leveraging MemorySeq tools, this is the first body of work that seeks to identify biomarkers for stress resistance in CHO using heritability principles. Ammonia stress has been shown to influence glycosylation, amino acid metabolism, and induce genomic instability in the form of indels and SNPs. Using bulk RNAseq to measure transcriptional shifts for ammonia-stressed CHO cells, genes relating to alanine metabolism, cell-cycle regulation, cellular senescence, and DNA damage/repair were identified as possibly contributing to impaired growth and productivity kinetics^35,40^. However, there was no metric for discerning the noise of differentially expressed genes and those that could play a key role in stress adaption. Alone, MemorySeq highlights gene states that play a crucial role in the transition to a stress tolerant state. However, not all genes may necessarily be responsible for the tolerance phenotype.rare^81^ By combining MemorySeq with stress adapted DEG, this method narrows the list down to possible entry points into stress response pathways. Similarly, exploratory works in hyperosmolar stress conditions characterized the same effect of enhanced specific productivity and reduced growth rate. Transcriptomic and proteomic analysis identified mitochondrial activation, oxidative stress reduction, and cell cycle progression as playing a role in hyperosmolar stress response, but could not comment further as to which genes were most important in the developing stress adapted phenotype^62,82^. Previous literature has widely considered the phenotypic and surface-level proteomic changes associated with manufacturing stress conditions but has not reflected on the possible pathways for stress adaptation.

There are many cell line engineering strategies available for targeting these biomarkers to induce a stress-resistant phenotype. If the activation of a gene is correlated with stress-resistance, gene overexpression could be achieved by integrating additional copies or using inactivated Cas9 (dCas9)-directed activation (CRISPRa) to enhance stress adaptation^83–85^. Likewise, if the repression of a gene is correlated with stress-resistance, gene knockdown-methods such as RNA interference (RNAi) or dCas9-directed inhibition (CRISPRi) could also be employed for the same effect^86,87^. However, stress response pathways are comprised of many proteins that influence a variety of molecular functions to accommodate and mitigate the deleterious effects of stress. Methods that target a single gene may not always activate a whole pathway but lessen only one symptom of the stress^88,89^. Activation of response pathways develops a more robust stress response to tackle the multi-faceted burden of cellular stress^90,91^. These stress-responsive pathways have been noted in CHO to alleviate oxidative-stress associated with production and bioreactor conditions that involve anti-apoptotic and GSH biosynthetic pathways^92–94^. In this work, 4 co-fluctuating community networks that contain genes relating to stress detection and critical cell health processes were identified and may demonstrate critical entry points into stress response pathways. It is believed that these heritable, but ultimately transient fluctuations in expression are largely due to epigenetic modifications and complex interactions or crosstalk between genes within network communities that display the co-fluctuations. While no comparable study has been conducted in CHO, it has been demonstrated in other organisms that temperature, nutrient, and osmolality induced environmental stress has resulted in epigenetically driven and inherited stress tolerant states. In *S. cerevisiae* during periods of inositol starvation and osmotic stress, accumulation of di- and tri-methylation of H3K4 histone marks was observed following a period of hyperacetylation of H3 and H4^95,96^. This state correlated to the recruitment of the SET3C complex that reinforced an active heritable memory that facilitated the recruitment of poised RNAPII lasting 4-8 generations^96^. A similar pattern of H3K4me2 deposition and transgenerational inheritance of stress resistance has been observed in eukaryotic organisms such as HeLa and *D. melanogaster* cells as facilitated by the nuclear pore protein Nup98.^97–99^ Additional epigenome characterization work must be conducted to verify this phenomenon in CHO. The network communities of co-fluctuating genes identified in this work are unlikely a reflection of genetic mutations, but rather influenced by epigenetics, chromatin accessibility, and ability of transcription factors to bind given the transiency of these rare phenotypes proven in the original MemorySeq work^26,60^. Regulating gene expression through conventional means neglects the positional and epigenetic context for these genes, possibly a vital component for the mechanism of co-fluctuation and activation of many genes in a pathway. A promising alternative comes in the form of site-specific epigenetic modifications to rewrite the epigenetic marks and modulate local chromatin structure. Localization of these marks at biomarkers is an alternative for regulating gene expression in a way that is heritable and taps into the higher order structure of chromosomal DNA. Doing so influences spatially adjacent genes that may synergize within stress responsive pathways. While the mechanism for the intermediate heritable states described in this worked are still ambiguous, the rapid and transient fluctuations suggest that epigenetics, which are also transient in nature, play a significant role. These states also appear to be a combination of conserved and cell line specific processes. When compared to the original MemorySeq work, performed in the WM989 and WM983B melanoma, MDA breast cancer, and PC9 carcinoma cell lines, any from 4% to 10% of the heritable gene states unique to these cell lines appeared in our CHO cell line. Across all four of the cell lines, 16.5% of the genes there were identified as heritable in this work appeared heritable in at least one of the other cell lines, including *Hmox1, Ier3,* and *Serpine1*^26^. These shared gene states may represent some form of evolutionarily conserved states that serve as important entry points to the transition to stress tolerance.

## 4. Conclusion

This study sought to expand our understanding of CHO cell stress adaptation. The deleterious phenotypic effects of common manufacturing stressors, such as ammonia, lactate, and osmolality, observed in literature were likewise seen here. However, using the novel transcriptomic workflow described in MemorySeq, the process for parsing the transcriptional noise that the stresses induce becomes more meaningful. The combination of information obtained from stress-induced cultures and MemorySeq reinforces our understanding of manufacturing stress while highlighting diagnostic genes for the rational design of engineered cell lines. While the exact mechanism of the bimodal systems observed here remains unclear, their observation in other systems reinforces the idea that epigenetics and, to some extent, stochasticity play an important role. Such epigenetically driven, bistable, long-term memory states have been elucidated in epithelial tissue and *S. enterica* to adapt to perturbations within the environment^19,20^. Being able to harness what is otherwise a natural and random adaptation phenomenon into an intentionally engineered pathway will lead to CHO strain that is primed for the strain of manufacturing-scale condition. Healthier cells remain more productive for longer and alleviate the burden of downstream purification. This pipeline for assessing stress-dependent effects is also translatable to other cell lines and other stress conditions, expanding our ability to identify cell line engineering targets and streamline CLD processes.

## 5. Acknowledgments and Funding

We thank the Expression Systems and Novel Biopharm Materials Team at MilliporeSigma for access to the CHOZN^®^ GS^-/-^ Clone 23 cell line; Dyllan Rives for aiding in cell culture; Molly Wintenberg for providing RNAseq analysis pipeline for CHO, Sylvain Le Marchand with the Delaware Biotechnology Institute’s Bio-Imaging Center for instruction and experimental planning for smRNA-FISH; Erin Bernberg and Mark Shaw with the Delaware Biotechnology Institute’s Sequencing and Genotyping for aid in RNA extractions and quality verification; Ross Klauer, Emily Doleh, Christopher Pirner for reviewing and editing the paper. This work was supported by funding from the UD College of Engineering.

## 6. Author Contributions

Conceptualization, S.G., A.S., and M.B.; Methodology, S.G., A.S., and M.B.; Software, S.G. and Z.D.; Formal Analysis S.G., A.S., and Z.D.; Investigation, S.G. and Z.D.; Visualization, S.G. and Z.D.; Data Curation, S.G.; Writing – Original Draft, S.G; Writing – Review and Editing, S.G., A.S., and M.B.; Resources, M.B., Supervision, M.B.; Funding Acquisition, M.B.;

## 7. Declaration of Interests

The authors declare no competing interests.

## 8. STAR Methods

### 8.1 Cell-culture and Maintenance

A recombinant CHOZN^®^ GS^-/-^ ZFN, a CHO-K1 subclone with glutamine synthetase (GS) knocked out for glutamine selection, provided by MilliporeSigma was used for this work. This subclone, referred to as CHOZN^®^ GS^-/-^ Clone 23, expressed a monoclonal antibody and GS for selection. These cells were grown in EX-CELL^®^ CD CHO Fusion Medium (Sigma-Aldrich, St. Louis, MO), deficient in glutamine and seeded and passaged in 125-mL Erlenmeyer flasks (Corning, Corning, NY) with a working volume of 25 mL. The cells were cultured in a Minitron incubator (INFORS HT, Bottmingen, Switzerland) at 37 °C, 5% CO_2_, 80% relative humidity, and shaking at 100 rpm with passages every 3 days at a cell density of 0.5×10^6^ cells/mL.

### 8.2 Single Cell Limited Dilution Cloning

Single cell limited dilution cloning (LDC) was used to isolate monoclonal pools for MemorySeq. To assist outgrowth with vital growth factors and nutrients during LDC, conditioned media was produced by harvesting sufficient cells in a 25 mL flask to achieve a seeding density of 1.0 x 10^6^ cells/mL. The cell culture media was harvested 24 hours post-initiation by centrifuging the culture medium at 200 x g for 5 minutes and sterile filtering the media through a 0.22 *μ*m syringe filter. Conditioned media was stored at 4 °C for no longer than seven days.

24 hours before starting LDC, a stock culture was seeded at 1.0 x 10^6^ cells/mL. Sufficient cell culture volume was collected and serially diluted to obtain a final concentration of 2.5 cells/mL in an 80%/20% mix of EX-CELL^®^ CHO Cloning Media (Sigma-Aldrich, St. Louis, MO) and conditioned media. Six non-treated 96-well plates (CELLTREAT, Pepperell, MA) were seeded with 200 *μ*L of the mix for an average of 0.5 cells/well. Plates incubated under static conditions at 37 °C, 5% CO_2_, and 80% relative humidity and were undisturbed for 7 days. Afterwards, plates were inspected using a light optical microscope and wells with outgrowth originating from a single location were noted as single cell pools while those with no outgrowth or outgrowth from multiple places were discarded. The plates were fed an additional 20 *μ*L of EX-CELL^®^ CHO Cloning Media to supplement growth and maintain volume after evaporation. After 3 weeks, single cell colonies reached roughly 100,000 cells or 80% confluency.

### 8.3 RNA Extraction and Sequencing

Cell samples were collected either from LDC plates, standard passage flasks, or from fed-batch flasks. For MemorySeq RNA extraction, 8 x 10^4^ – 12 x 10^4^ cells were collected for extraction and for fed-batch flasks, 2.5 x 10^6^ cells were used for extraction. RNA was extracted using the miRNeasy Mini Kit (Qiagen, Germantown, MD) following manufacturer’s protocol. All RNA samples were run on the Agilent 5200 Fragment Analyzer (Agilent, Santa Clara, CA) to verify RNA quality prior to submission for sequencing. Samples with an RIN ≥ 6.0 and at least 100 ng of high quality RNA were sent for sequencing. For MemorySeq, the target was 40 RNAseq samples for both MemorySeq monoclonal samples and noise control samples, but after quality selection only 38 samples and 40 samples remained for MemorySeq samples and noise control samples respectively. RNAseq samples were submitted to Azenta for RNA selection using poly(A) selection and library preparation. Samples were sequenced on the Illumina^®^ HiSeq 4000^®^ and were sequenced at a coverage of at least 19x and on average 24x with at least uniquely mapped 10 million reads per RNA sequencing library. Sequencing files were returned as FASTQ files and stored online at GEO Series accession number GSE232813 (https://www.ncbi.nlm.nih.gov/geo/query/acc.cgi?acc=GSE232813) along with the converted count tables. Raw FASTQ files were transferred to the University of Delaware Biomix HPC cluster where they were processed through a pipeline consisting of Trim Galore for adapter trimming/quality control, STAR for RNAseq alignment to CriGri-PICR, Samtools for file conversion, and HTSeq to enumerate unique and high-quality mapped reads^100^. Gene count tables were transferred to RStudio and processed further depending on the application. The RNAseq processing pipeline is available at https://github.com/SGrissomUDel/CHOCell_MemorySeq.

### 8.4 MemorySeq Processing

For the computational analysis of MemorySeq samples and noise control samples, the raw count for each gene was converted to transcripts per million (TPM). The genes were filtered out such that only protein-coding genes and TPM ≥2.5 were included in the dataset. Metrics of variation, including average, standard deviation, coefficient of variation, skewness, and kurtosis were all calculated for each gene in each dataset. It was found that the relationship between log_2_(TPM) and coefficient of variation for most genes was well-described by a Poisson regression model. Heritable gene expressions states in MemorySeq sample were observed to deviate from this relationship. As such, heritable gene expression states were defined as genes with TPM ≥2.5 and in the 98^th^ percentile for residuals from the Poisson regression fit. Any gene expression states in the noise control that exceeded the residual cutoff were removed from the heritable pool. This yielded 199 unique heritable gene expression states. To understand how heritable gene expression states co-fluctuate with each other, a correlation matrix was generated using pairwise Pearson correlation coefficients. The resulting matrix, containing all unique significantly heritable gene expression states, had dimensions of 199 genes x 199 genes. Computational analysis and MemorySeq processing tools are available at https://github.com/SGrissomUDel/CHOCell_MemorySeq (See **Supplemental File 1** and **Supplemental File 2**).

Using the Pearson correlation coefficients matrix for all significantly heritable gene expression state, network communities were generated using a *k*-clique percolation method (*k*- CPM) using the CliquePercolation (0.3.0) package in R. This algorithm detects communities by assigning each gene as an independent node and each node is connected to each other by edges defined as the correlation in the undirected and weighted Pearson correlation matrix. There are two adjustable parameters when generating the network communities. The first is *I*, which defines the intensity threshold or the absolute value of the edge threshold required to connect two nodes. The other is *k*, which defines the minimum number of fully connected nodes to form a *k*- clique. A *k*-clique community then contains all adjacent *k*-cliques that share *k*-1 nodes. A value of *k*=4 was used based on prior *k*-CPM for similar analyses while *I* was varied from 0.6 to 0.98 in order to find the optimal threshold^54,55^. Adjusting *I* changes the size of each *k*-clique community, the number of communities, and the number of isolated nodes. Optimization of *I* can be achieved by tuning *I* such that the ratio of the largest to the second largest community is roughly two and/or maximizing variance after excluding the largest community, where variance, *χ*, is defined as shown in ***Equation 1*** where *N* is the total number of communities detected, *n_i_* is the size of the *i^th^*community excluding the largest one, and *n*_$_ is the size of the *j^th^*community excluding the largest community and the *i^th^* community.

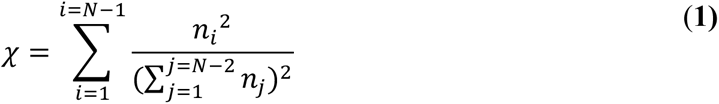

Gene ontology (GO) enrichment analysis, a method used to detect the over-representation of certain gene product attributes within a group of genes, was used to assign enriched biological processes to each community to further understand the functional similarities of the co-fluctuating gene networks. GO biological processes terms were collected for 13,288 of the 21,386 protein-coding genes for the CriGri-PICR assembly^100^. GO enrichment analysis was completed using the topGO (2.46.0) R package with a minimum node size of 10. The statistical test of significance was conducted using Fisher’s exact test with the classic, weighted, and elimination methods to ensure physically significant enrichment was captured with an adjusted p value < 0.05 (See **<u>Supplemental File 4</u>**: GO Enrichment Analysis).

### 8.5 Prediction of Heritability Lifetime Index

Expression states were modeled as either inheriting the bulk average phenotype (OFF) or a rare deviating phenotype (ON). Exponential cell proliferation or growth rate was modelled by *k_x_* (a generation time of 1/*k_x_*). MemorySeq clonally derived pools were assumed to be in a constant state of exponential growth throughout the duration of the experiment, (from one cell to divide into 100,000 cells), with a consistent proliferation rate *k_x_*. This proliferation rate was also assumed to be constant with respect to whether an indivudal cell was in the ON or OFF state. The rate of fluctuation or transition from the ON to OFF state was denoted as *k_OFF_* while the inverse transition was denoted as *k_ON_*. The fraction of cells displaying the rare ON state in the bulk population, *f*, was defined in ***Equation 2***, or as the ratio of rates of transition. Previous work conducted by Shaffer et. al. found *f* was typically 1% or less in RNA-FISH and internally confirmed by flow-cytometry^26^.

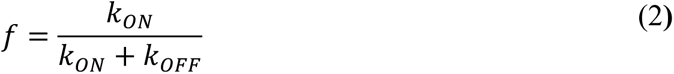

In this fluctuation analysis, any given cell sampled from the bulk population has a probability *f* of existing in the rare ON state and probability 1-*f* of existing in the OFF state. The random variables *x(t)* and *y(t)* are then defined as the total number of cells in a given sample and time *t* in the ON and OFF state respectively. The stochastic time evolution of *x(t)* and *y(t)* is dictated by proliferation and the generation of new cells that inherit the ON or OFF state and fluctuations by existing cells between the ON and OFF state. The ratio *x(t)/(x(t) + y(t))* then represents the fraction of cells in the ON state at a given time *t*. Using the assumption that *f* << 1 and the first two statistical moments of *x(t)* and *y(t)*, derived using moment dynamics of stochastic systems (see Singh and Hespanha), the *CV^2^* of the ratio *x(t)/(x(t) + y(t))* is given in ***Equation 3***, where *T = tk*_*x*_, represents the duration of the experiment normalized to generation count (*T* ≈17 generations) and *T_ON_* = *t k_x_/k_OFF_* represents the average duration of the ON state normalized to generation count^101,102^.

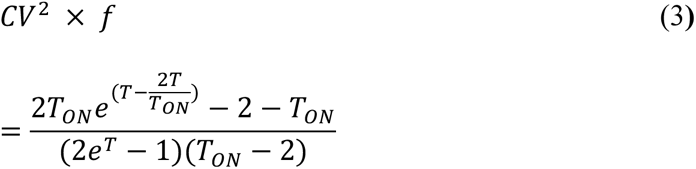

Application of *a priori* knowledge of *f* by assuming its value to be 0.01, an inverse transformation of ***Equation 3*** can be used to estimate *T_ON_* and the lifetime of certain heritable gene states (See **Supplemental File 3**: Heritability Index for Gene States Identified as Heritable)

### 8.6 smRNA-FISH and Imaging

Small molecule RNA fluorescence *in-situ* hybridization (smRNA-FISH) was used to visualize the relationship between cells sharing a common lineage and the maintenance of heritable gene expression states. RNA FISH probes were designed using the Stellaris^®^ Probe Designer software (LGC, Biosearch Technologies, Hoddesdon, United Kingdom) and included 30-48 probes that had length a 20 bp, at least 2 bp of spacing, and either Quasar^®^ 570 or Quasar^®^ 670 fluorescent dyes. These probes were designed to target Hmox1 and Ier3 as identified from MemorySeq. To track the shared lineage of dividing cells, 10,000 CHOZN^®^ GS^-/-^ Clone 23 cells suspended in 90% EX-CELL^®^ CD CHO Fusion Medium/10% FBS were first fixed onto an 18 mm diameter, #1 thickness fibronectin coated coverglass by centrifugation with 30 minute incubation at 37 °C. The coverglasses were incubated at statically at 37 °C, 5% CO_2_, and 80% relative humidity for 7 days or until 50-60% confluency after unattached cells were gently washed off.

Multiplex smRNA-FISH was achieved using the Stellaris^®^ RNA FISH reagents (LGC, Biosearch Technologies, Hoddesdon, United Kingdom) according to manufacturer’s protocol for adherent cells. Briefly, this involved fixation using 3.7% v/v formaldehyde solution in 1x PBS for 10 minutes and permeabilization using 70% ethanol for storage at 4°C. Before hybridization, coverglasses were washed with a wash buffer containing 10% formamide. Coverglasses were then transferred to a humidity chamber and incubated in a hybridization buffer containing 10% formamide and a Quasar^®^ 570 or Quasar^®^ 670 labeled RNA FISH probe and incubated for 16 hours. After hybridization, coverglasses were washed with a 10% formamide wash buffer before staining with DAPI. After mounting with Vectashield Mounting Medium, samples were imaged immediately using a Stellaris 8 tauSTED/FLIM confocal microscope (Leica Microsystems, Wetzlar, Germany).

### 8.7 Stress-Condition Fed-Batch and Analysis

Fed-batch flasks were established to adapt the CHOZN^®^ GS^-/-^ Clone 23 cell line to stress agents characteristic of manufacturing conditions. Fed-batch flasks were seeded from freshly thawed cells after 3 passages to allow recovery. The flasks were seeded at a cell density of 0.5 x 10^6^ cells/mL in 30 mL of EX-CELL^®^ Advanced CHO Fed-batch Medium (Sigma-Aldrich, St. Louis, MO). Starting day 3, EX-CELL^®^ Advanced CHO Feed 1 (Sigma-Aldrich, St. Louis, MO) was fed at 5% of the total volume every other day and D-(+)-glucose solution was fed to maintain a concentration of 4 g/L of glucose starting day 4. Cell viability and cell density was measured daily using the DeNovix^®^ CellDrop (DeNovix, Wilmington, DE). Supernatant and cell pellets were collected every other day starting day 1 and cell samples were stored in RNAlater stabilization solution (ThermoFisher Scientific, Waltham, MA) after washing with 1x PBS. Fed-batch was carried out until all flasks fell below 70% cell viability. Ammonium chloride (VWR, Radnor, PA), sodium lactate (Sigma-Aldrich, St. Louis, MO), and sodium chloride (Sigma-Aldrich, St. Louis, MO) were all spiked in at the beginning of fed-batch at varying concentrations to simulate stress during fed-batch production.

Transcriptomic analysis was conducted on cell-samples by first extracting RNA, submitting for sequencing, and processing the results as described earlier. They were sequenced at a coverage of at least 65x and on average 75x with at least 35 million uniquely mapped reads per RNA sequencing library. Gene count tables were transferred to RStudio and differential gene expression analysis (DGEA) was conducted using the DESeq2 (1.34.0) package. A log_2_ fold change (L2FC) threshold of 0.58 or a 1.5-fold change and an adjusted p-value of 0.05 defined genes with significant differential expression. The intersection of differentially expressed genes and heritable gene expression states was used in GO enrichment analysis to find significantly overrepresented biological processes (See **<u>Supplemental File 4</u>**: GO Enrichment Analysis).

## 11. Supplemental Figures

**Figure S1:**
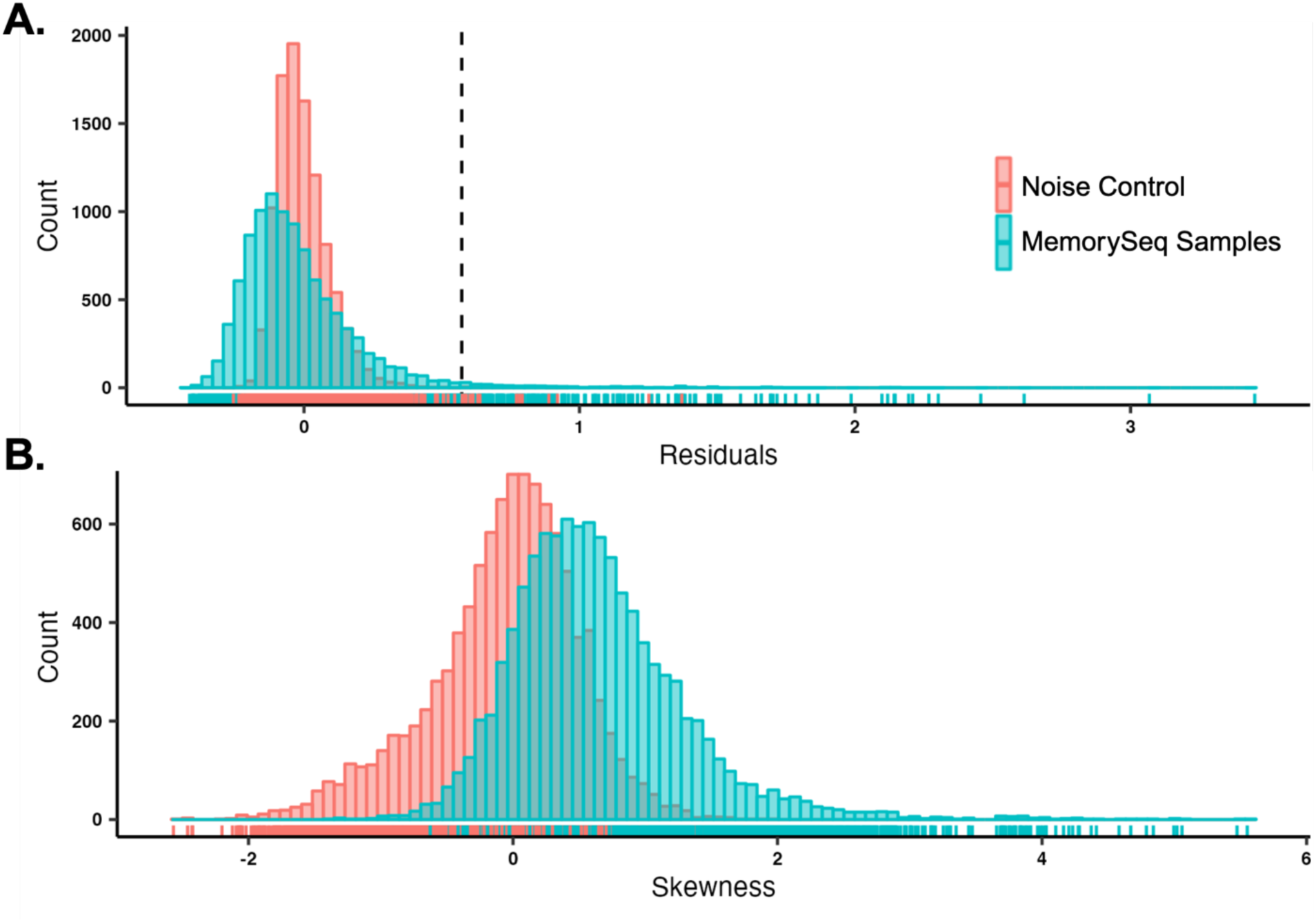
Visualization of Variability and Skew for MemorySeq Clones: (A.) Distribution for 10,106 protein coding genes for their residuals relative the fitted Poisson regression model. The dotted line represents the residual threshold, above which genes were considered to display a heritable pattern. (B.) Distribution of skew for all genes, where the MemorySeq samples are shifted to the right, a reflection of greater variability from single-cell derived parents rather than bulk

**Figure S2:**
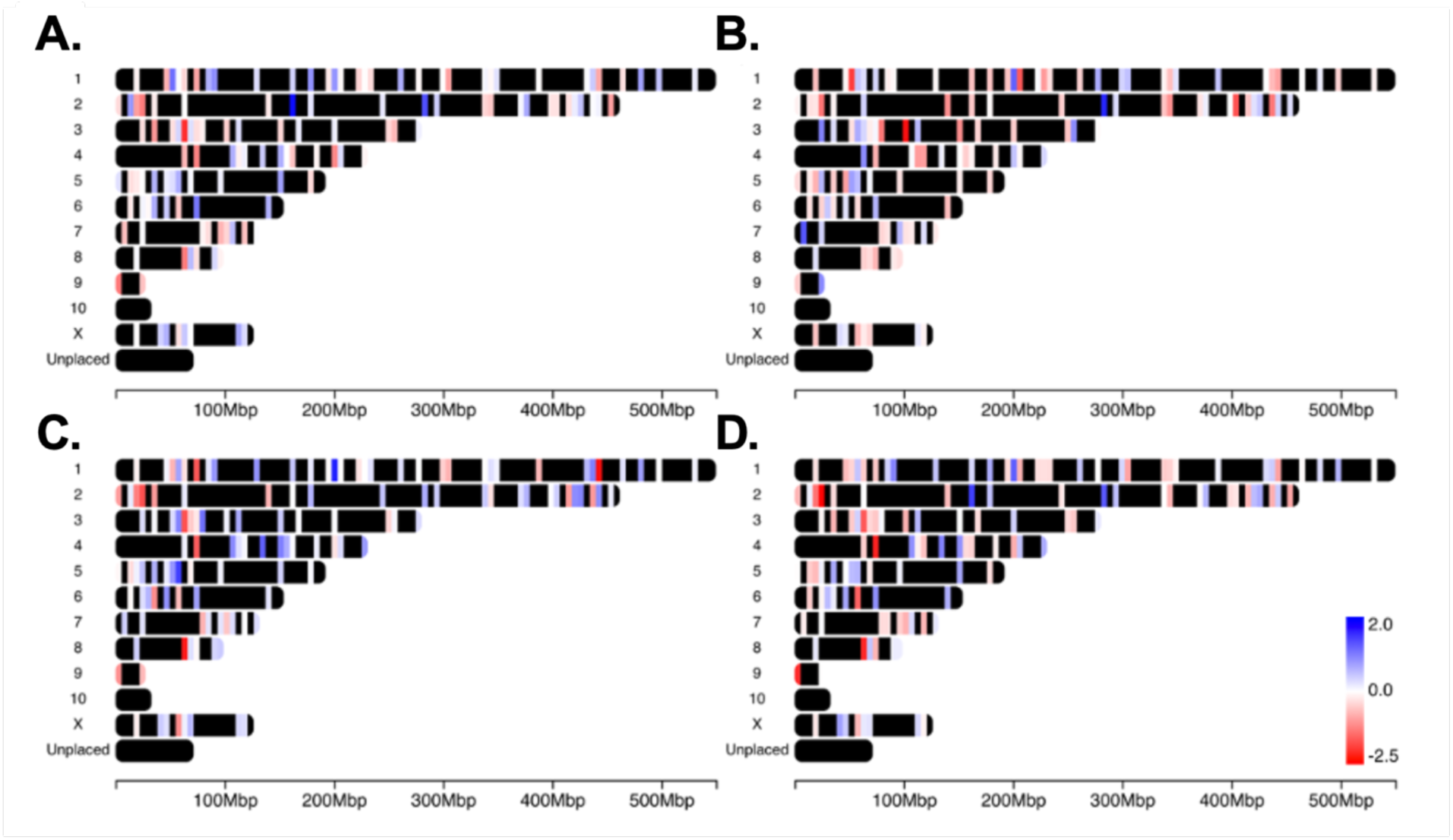
Mapping of genes with heritable expression patterns reveals even spatial distribution throughout genome: Spatial distribution of heritable genes compared to the CriGri PICR genome assembly. The colors reflect the log_2_fold change in expression for the fed-batch flasks under (A.) high ammonia stress, (B.) high lactate, (C.) high osmolality, and (D.) high combination stress.

**Figure S3:**
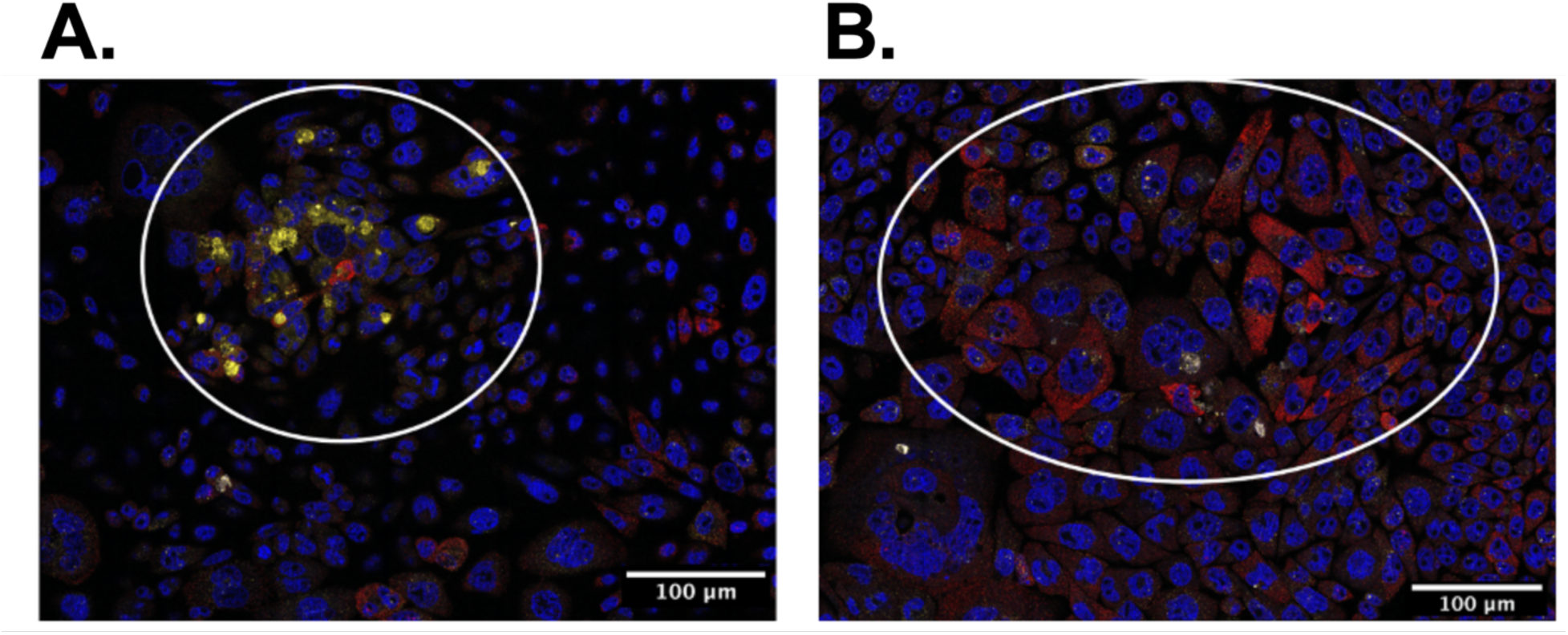
Mapping of genes with heritable expression patterns reveals even spatial distribution throughout genome: RNA FISH image collected for CHOZN^®^ GS^-/-^ Clone 23 cells fixed on a fibronectin surface after one week of expansion. Cells were stained with DAPI (shown in blue) and multiplexed with a Quasar® 670 labelled Hmox1 FISH probe (shown in red) and a Quasar® 570 labelled Ier3 FISH probe (shown in yellow). Pockets of highly expressing cells provide evidence of heritability of rare expression states as proximity is directly correlated with relatedness. (A). Highlights a region of high Ier3 expression and (B.) highlights a region of high Hmox1 expression.

**Figure S4:**
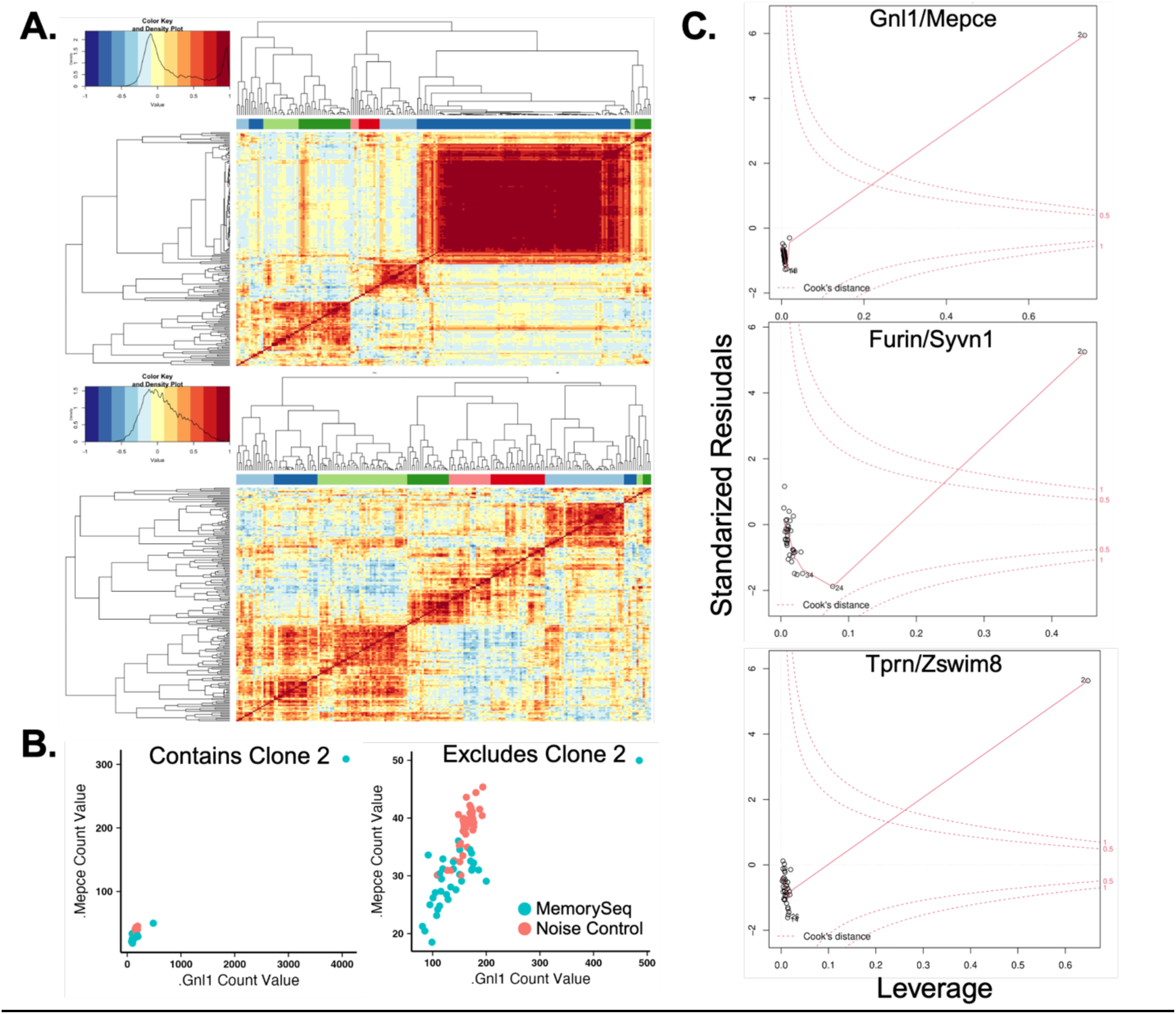
smRNA-FISH captures highly expressing clones in close proximity which serves as an estimator for relatedness: Characterization of outliers within the MemorySeq Clones. (A.) Pearson pairwise correlation matrix for all 199 heritable gene states with (Top) and without (Bottom) the MemorySeq Clone 2 that was later identified as an outlier. Distribution of correlations show a sharp increase of correlations in the 0.9-1.0 range with Clone 2 included. The exclusion of Clone 2 shows a more Gaussian distribution of correlations (B.) Representative scatterplots of a strongly correlating gene pair, Mepce and Gnl1. The inclusion of Clone 2 shows a significant outlier biasing the correlation strength. The exclusion highlights a positive, yet significantly weaker correlation. (C.) Cook’s distance plots for three gene pairs with correlations in the 0.95 to 1.0 range. The plot visualizes the standardized residuals versus the leverage for each of the 38 MemorySeq clones. The leverage is a measurement of how influential each clone is on establishing the correlation value. The red dotted lines mark a Cook’s distance greater than 0.5 and 1. A Cook’s distance greater than 1 indicates a point as an outlier. While outliers are intrinsic to this dataset, clone 2 proved to dominate the correlation values and confounded the variability the other 37 clones contributed.

**Figure S5:**
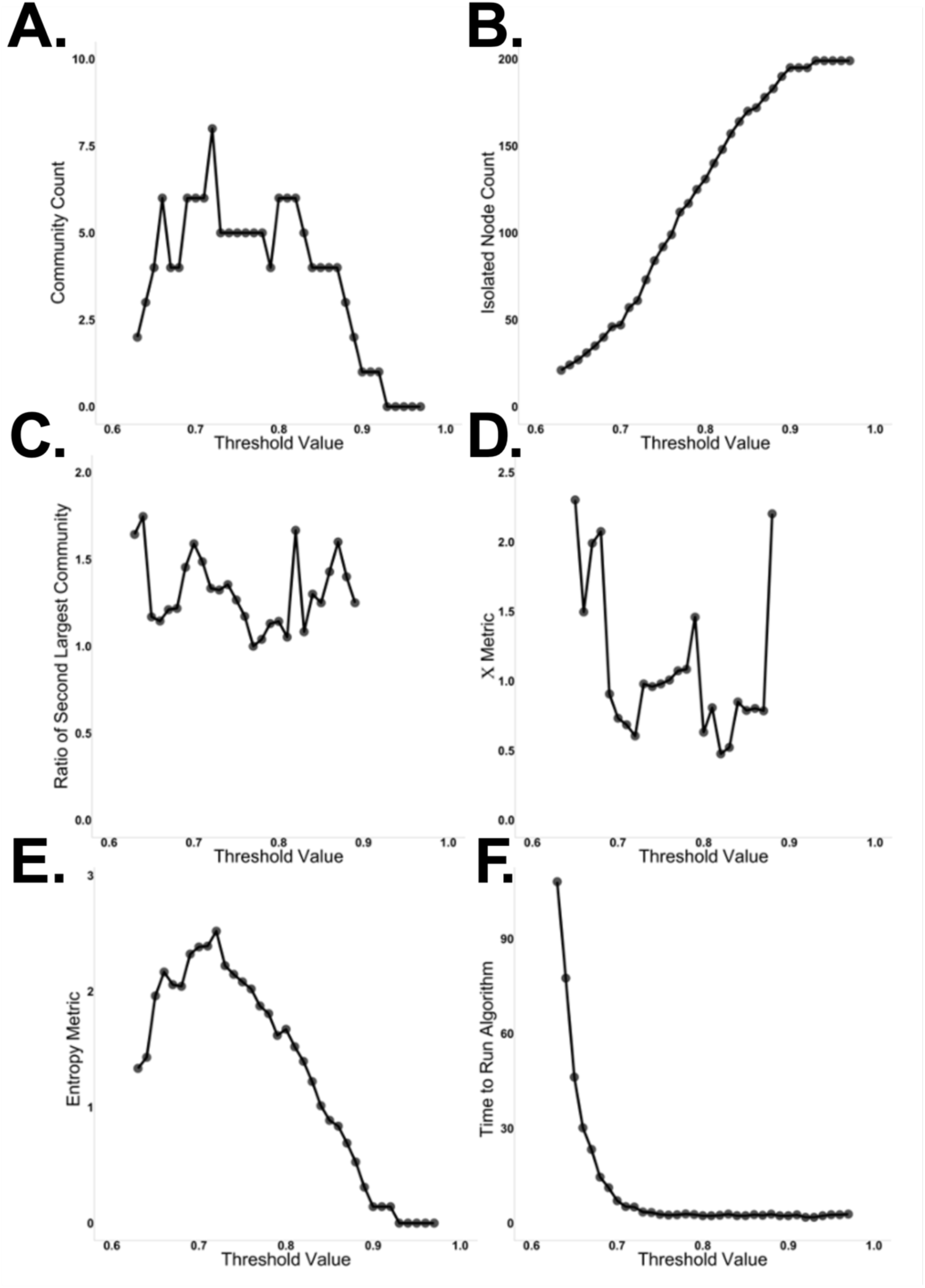
Statistical heuristic for identifying critical threshold parameter for k-clique percolation for network community identification: Optimization process for determining threshold value for k-clique percolation for network community identification. The chosen value of I*t*0.68 was chosen given a balance of a large entropy metric, a large *χ* metric, and a ratio of largest to second largest community was in the range of 1.0 to 2.0. (A.) Number of distinct network communities identified given k*t*4 across varying threshold values. (B.) Number of isolated nodes, or genes, across varying threshold values. (C.) Ratio of largest to second largest community, ideal being ∼2.0, versus threshold values, (D.) Variance, or *χ* metric, after excluding the largest community versus threshold values and should ideally be maximized. (E.) Entropy metric versus threshold values and should ideally be maximized. (F.) Time to run the percolation algorithm in minutes versus threshold values. Shows the complexity of lower threshold systems and the exponential increase in community structure.

**Figure S6:**
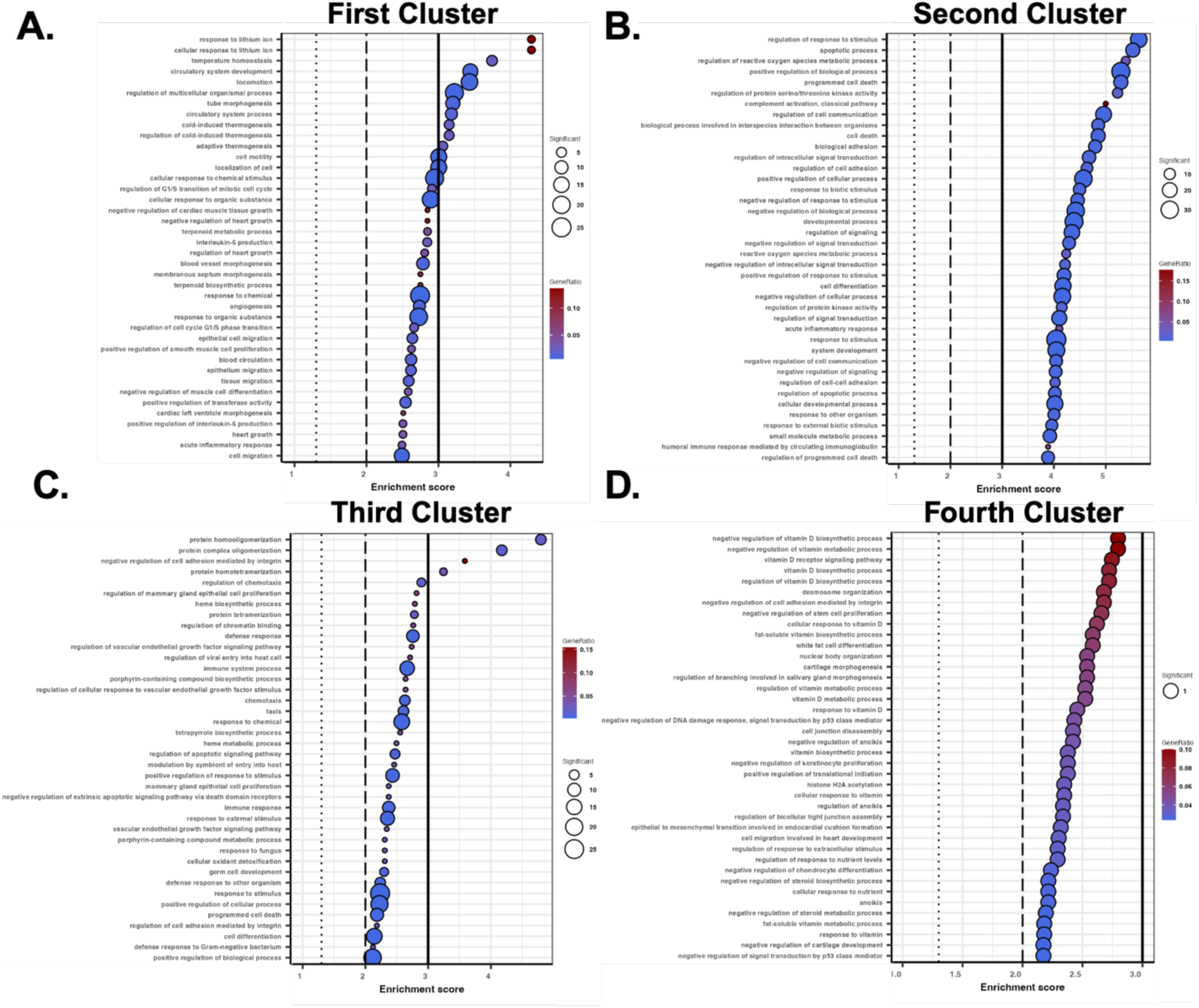
Classic Fisher GO enrichment analysis for heritable gene clusters: Dot plot visualizing the 40 most enriched GO terms across all 199 heritable gene states. Size and color intensity reflect the number of genes contained with each GO term and the corresponding gene ratio. Dotted lines represent a p-value of 0.05, 0.01, and 0.001 from left to right. (A.) First Cluster (red) identified from the community network, (B.) Second Cluster (green) identified from the community network, (C.) Third Cluster (teal) identified from the community network, and (D.) Fourth Cluster (purple) identified from the community network.

**Figure S7:**
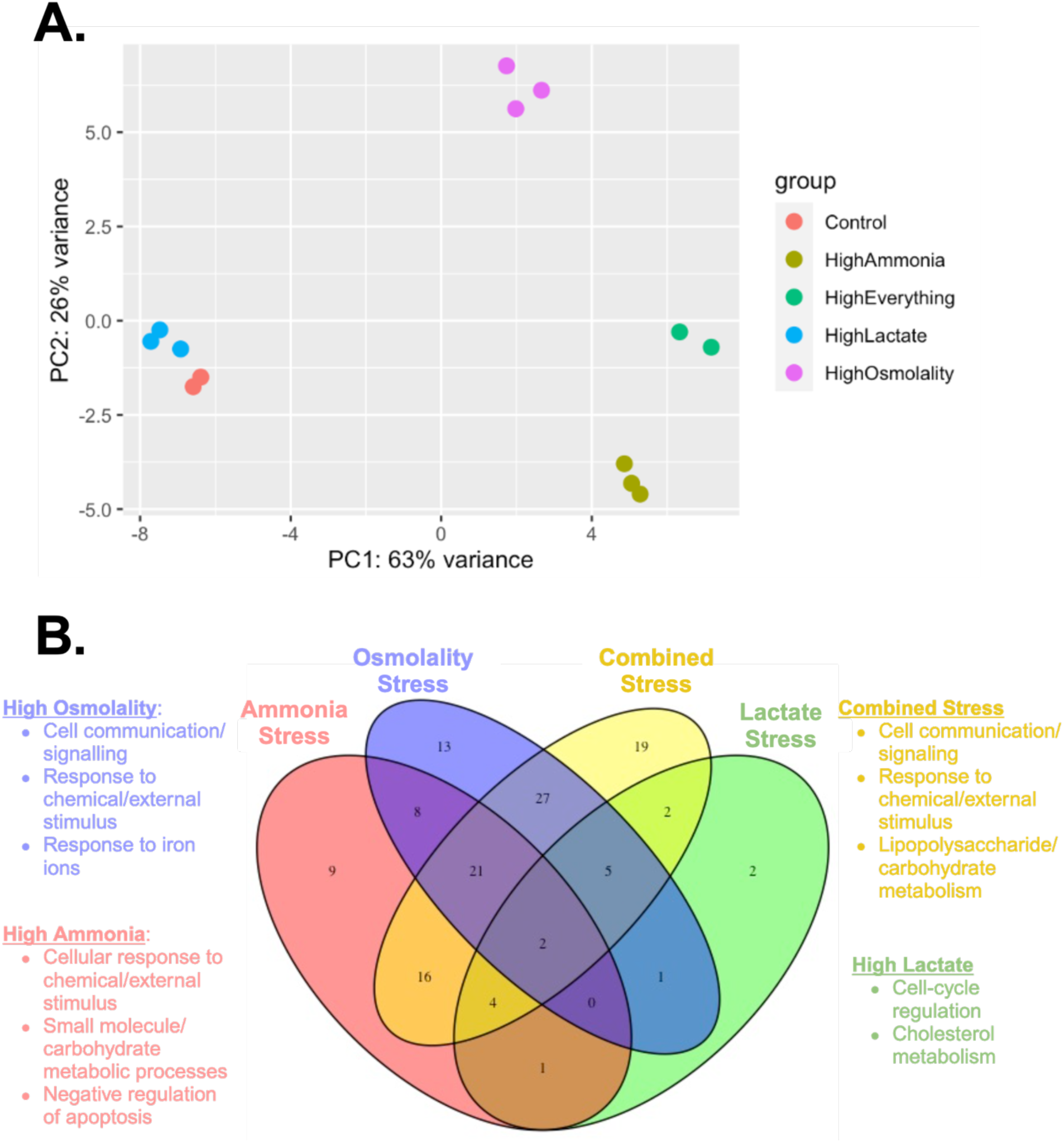
Statistical characterization of different stress conditions reveals distinctly unique differential expression patterns: (A.) Principle component analysis assessing the transcriptomic variability amongst the different stress conditions. Spatial distance along the PC1 direction marks a more significant difference than spatial distance along the PC2 direction. Thus, there was significant transcriptomic variability between high ammonia, osmolality, and combination and the control flask while there was not a significant difference between high lactate and the control. (B.) Overlap of DEG and heritable gene states identified using MemorySeq and commonalities between different manufacturing stressors. GO enrichment analysis was carried out to see what biological processes were overrepresented in the DEG

## 12. Supplemental Files

**Supplemental File 1**: Coefficient of Variance and Summary Statistics for Gene States Identified as Heritable in Noise Control Samples

CV_For_HeritableGenes_Noise_Controls.xlsx

**Supplemental File 2**: Coefficient of Variance and Summary Statistics for Gene States Identified as Heritable in MemorySeq Samples

CV_For_HeritableGenes_MemorySeq_Samples.xlsx

**Supplemental File 3**: Heritability Index for Gene States Identified as Heritable

HeritabilityIndex.xlsx

**Supplemental File 4**: GO Enrichment Analysis

GO_EnrichmentAnalysis.xlsx

## Notes

### Competing Interest Statement

The authors have declared no competing interest.

## Works Referenced

1. Wildt, S. & Gerngross, T. U. The humanization of N-glycosylation pathways in yeast. Nat. Rev. Microbiol. 3, 119–128 (2005).

2. Brooks, S. A. Appropriate glycosylation of recombinant proteins for human use. Mol. Biotechnol. 28, 241–255 (2004).

3. Lee, J. S., Kildegaard, H. F., Lewis, N. E. & Lee, G. M. Mitigating Clonal Variation in Recombinant Mammalian Cell Lines. Trends Biotechnol. 37, 931–942 (2019).

4. Kim, M., O’Callaghan, P. M., Droms, K. A. & James, D. C. A mechanistic understanding of production instability in CHO cell lines expressing recombinant monoclonal antibodies. Biotechnol. Bioeng. 108, 2434–2446 (2011).

5. Ko, P. et al. Probing the importance of clonality: Single cell subcloning of clonally derived CHO cell lines yields widely diverse clones differing in growth, productivity, and product quality. Biotechnol. Prog. 34, 624–634 (2018).

6. Baik, J. Y. & Lee, K. H. Growth Rate Changes in CHO Host Cells Are Associated with Karyotypic Heterogeneity. Biotechnol. J. 13, 1700230 (2018).

7. Spahn, P. N. et al. Restoration of DNA repair mitigates genome instability and increases productivity of Chinese hamster ovary cells. Biotechnol. Bioeng. 119, 963–982 (2022).

8. Keung, A. J., Joung, J. K., Khalil, A. S. & Collins, J. J. Chromatin regulation at the frontier of synthetic biology. Nat. Rev. Genet. 16, 159–171 (2015).

9. Park, M., Keung, A. J. & Khalil, A. S. The epigenome: the next substrate for engineering. Genome Biol. 17, 183 (2016).

10. Turner, B. M. Epigenetic responses to environmental change and their evolutionary implications. Philos. Trans. R. Soc. B Biol. Sci. 364, 3403–3418 (2009).

11. Feichtinger, J. et al. Comprehensive genome and epigenome characterization of CHO cells in response to evolutionary pressures and over time. Biotechnol. Bioeng. 113, 2241–2253 (2016).

12. Veith, N., Ziehr, H., MacLeod, R. A. F. & Reamon-Buettner, S. M. Mechanisms underlying epigenetic and transcriptional heterogeneity in Chinese hamster ovary (CHO) cell lines. BMC Biotechnol. 16, 6 (2016).

13. Hernandez, I. et al. Epigenetic regulation of gene expression in Chinese Hamster Ovary cells in response to the changing environment of a batch culture. Biotechnol. Bioeng. 116, 677–692 (2019).

14. Bintu, L. et al. Dynamics of epigenetic regulation at the single-cell level. Science 351, 720–724 (2016).

15. Lensch, S. et al. Dynamic spreading of chromatin-mediated gene silencing and reactivation between neighboring genes in single cells. eLife 11, e75115 (2022).

16. Lempiäinen, J. K. & Garcia, B. A. Characterizing crosstalk in epigenetic signaling to understand disease physiology. Biochem. J. 480, 57–85 (2023).

17. Suganuma, T. & Workman, J. L. Crosstalk among Histone Modifications. Cell 135, 604– 607 (2008).

18. Amabile, A. et al. Inheritable Silencing of Endogenous Genes by Hit-and-Run Targeted Epigenetic Editing. Cell 167, 219–232.e14 (2016).

19. Fernández-Fernández, R. et al. Evolution of a bistable genetic system in fluctuating and nonfluctuating environments. Proc. Natl. Acad. Sci. 121, e2322371121 (2024).

20. Clark, H. R. et al. Epigenetically regulated digital signaling defines epithelial innate immunity at the tissue level. Nat. Commun. 12, 1836 (2021).

21. Schuh, L. et al. Gene networks with transcriptional bursting recapitulate rare transient coordinated high expression states in cancer. Cell Syst. 10, 363–378.e12 (2020).

22. Gómez-Schiavon, M. & Buchler, N. E. Epigenetic switching as a strategy for quick adaptation while attenuating biochemical noise. PLoS Comput. Biol. 15, e1007364 (2019).

23. Park, J., et al. Epigenetic switch from repressive to permissive chromatin in response to cold stress. Proc. Natl. Acad. Sci. 115, E5400–E5409 (2018).

24. Walter, M., Teissandier, A., Pérez-Palacios, R. & Bourc’his, D. An epigenetic switch ensures transposon repression upon dynamic loss of DNA methylation in embryonic stem cells. eLife 5, e11418 (2016).

25. Ravindran Menon, D., Hammerlindl, H., Torrano, J., Schaider, H. & Fujita, M. Epigenetics and metabolism at the crossroads of stress-induced plasticity, stemness and therapeutic resistance in cancer. Theranostics 10, 6261–6277 (2020).

26. Shaffer, S. M. et al. Memory Sequencing Reveals Heritable Single-Cell Gene Expression Programs Associated with Distinct Cellular Behaviors. Cell 182, 947–959.e17 (2020).

27. Luria, S. E. & Delbrück, M. Mutations of Bacteria from Virus Sensitivity to Virus Resistance. Genetics 28, 491–511 (1943).

28. Chen, C., Liao, Y. & Peng, G. Connecting past and present: single-cell lineage tracing. Protein Cell 13, 790–807 (2022).

29. Dumont, J., Euwart, D., Mei, B., Estes, S. & Kshirsagar, R. Human cell lines for biopharmaceutical manufacturing: history, status, and future perspectives. Crit. Rev. Biotechnol. 36, 1110–1122 (2016).

30. Dahodwala, H. & Lee, K. H. The fickle CHO: a review of the causes, implications, and potential alleviation of the CHO cell line instability problem. Curr. Opin. Biotechnol. 60, 128–137 (2019).

31. Lai, T., Yang, Y. & Ng, S. K. Advances in Mammalian Cell Line Development Technologies for Recombinant Protein Production. Pharmaceuticals 6, 579–603 (2013).

32. Noh, S. M., Shin, S. & Lee, G. M. Comprehensive characterization of glutamine synthetase-mediated selection for the establishment of recombinant CHO cells producing monoclonal antibodies. Sci. Rep. 8, 5361 (2018).

33. Gomez, N. et al. Analysis of Tubespins as a suitable scale-down model of bioreactors for high cell density CHO cell culture. Biotechnol. Prog. 33, 490–499 (2017).

34. Lindgren, K. et al. Automation of cell line development. Cytotechnology 59, 1–10 (2009).

35. Chitwood, D. G. et al. Characterization of metabolic responses, genetic variations, and microsatellite instability in ammonia-stressed CHO cells grown in fed-batch cultures. BMC Biotechnol. 21, 4 (2021).

36. Schellenberg, J. et al. Stress-induced increase of monoclonal antibody production in CHO cells. Eng. Life Sci. 22, 427–436 (2022).

37. Shen, D. et al. Transcriptomic responses to sodium chloride-induced osmotic stress: A study of industrial fed-batch CHO cell cultures. Biotechnol. Prog. 26, 1104–1115 (2010).

38. Sieck, J. et al. Response of a CHO production cell line to different types of hydrodynamic shear stress present in stirred and sparged bioreactors. New Biotechnol. 29, S165 (2012).

39. Handlogten, M. W., Zhu, M. & Ahuja, S. Intracellular response of CHO cells to oxidative stress and its influence on metabolism and antibody production. Biochem. Eng. J. 133, 12–20 (2018).

40. Synoground, B. F. et al. Transient ammonia stress on Chinese hamster ovary (CHO) cells yield alterations to alanine metabolism and IgG glycosylation profiles. Biotechnol. J. 16, 2100098 (2021).

41. Li, J., Wong, C. L., Vijayasankaran, N., Hudson, T. & Amanullah, A. Feeding lactate for CHO cell culture processes: Impact on culture metabolism and performance. Biotechnol. Bioeng. 109, 1173–1186 (2012).

42. Slivac, I., Blajić, V., Radošević, K., Kniewald, Z. & Gaurina Srček, V. Influence of different ammonium, lactate and glutamine concentrations on CCO cell growth. Cytotechnology 62, 585–594 (2010).

43. Tauffenberger, A., Fiumelli, H., Almustafa, S. & Magistretti, P. J. Lactate and pyruvate promote oxidative stress resistance through hormetic ROS signaling. Cell Death Dis. 10, 1–16 (2019).

44. Zagari, F., Jordan, M., Stettler, M., Broly, H. & Wurm, F. M. Lactate metabolism shift in CHO cell culture: the role of mitochondrial oxidative activity. New Biotechnol. 30, 238–245 (2013).

45. Kamachi, Y. & Omasa, T. Development of hyper osmotic resistant CHO host cells for enhanced antibody production. J. Biosci. Bioeng. 125, 470–478 (2018).

46. Freund, N. W. & Croughan, M. S. A Simple Method to Reduce both Lactic Acid and Ammonium Production in Industrial Animal Cell Culture. Int. J. Mol. Sci. 19, 385 (2018).

47. Yoon, S. K., Choi, S. L., Song, J. Y. & Lee, G. M. Effect of culture pH on erythropoietin production by Chinese hamster ovary cells grown in suspension at 32.5 and 37.0 degrees C. Biotechnol. Bioeng. 89, 345–356 (2005).

48. Christie, A. & Butler, M. The adaptation of BHK cells to a non-ammoniagenic glutamate-based culture medium. Biotechnol. Bioeng. 64, 298–309 (1999).

49. Altamirano, C., Paredes, C., Cairó, J. J. & Gòdia, F. Improvement of CHO cell culture medium formulation: simultaneous substitution of glucose and glutamine. Biotechnol. Prog. 16, 69–75 (2000).

50. Maranga, L. & Goochee, C. F. Metabolism of PER.C6 cells cultivated under fed-batch conditions at low glucose and glutamine levels. Biotechnol. Bioeng. 94, 139–150 (2006).

51. Dhama, K. et al. Biomarkers in Stress Related Diseases/Disorders: Diagnostic, Prognostic, and Therapeutic Values. Front. Mol. Biosci. 6, 91 (2019).

52. Giusti, C., Pastalkova, E., Curto, C. & Itskov, V. Clique topology reveals intrinsic geometric structure in neural correlations. Proc. Natl. Acad. Sci. 112, 13455–13460 (2015).

53. Rieck, B., Fugacci, U., Lukasczyk, J. & Leitte, H. Clique Community Persistence: A Topological Visual Analysis Approach for Complex Networks. IEEE Trans. Vis. Comput. Graph. 24, 822–831 (2018).

54. Palla, G., Derényi, I., Farkas, I. & Vicsek, T. Uncovering the overlapping community structure of complex networks in nature and society. Nature 435, 814–818 (2005).

55. Jonsson, P. F., Cavanna, T., Zicha, D. & Bates, P. A. Cluster analysis of networks generated through homology: automatic identification of important protein communities involved in cancer metastasis. BMC Bioinformatics 7, 2 (2006).

56. Sha, S., Bhatia, H. & Yoon, S. An RNA-seq based transcriptomic investigation into the productivity and growth variants with Chinese hamster ovary cells. J. Biotechnol. 271, 37–46 (2018).

57. Fischer, S., Handrick, R. & Otte, K. The art of CHO cell engineering: A comprehensive retrospect and future perspectives. Biotechnol. Adv. 33, 1878–1896 (2015).

58. Fulda, S., Gorman, A. M., Hori, O. & Samali, A. Cellular Stress Responses: Cell Survival and Cell Death. Int. J. Cell Biol. 2010, 214074 (2010).

59. Nie, Z. et al. c-Myc is a universal amplifier of expressed genes in lymphocytes and embryonic stem cells. Cell 151, 68–79 (2012).

60. Sun, M. & Zhang, J. Chromosome-wide co-fluctuation of stochastic gene expression in mammalian cells. PLoS Genet. 15, e1008389 (2019).

61. Hilliard, W. & Lee, K. H. Systematic identification of safe harbor regions in the CHO genome through a comprehensive epigenome analysis. Biotechnol. Bioeng. 118, 659–675 (2021).

62. Alhuthali, S., Kotidis, P. & Kontoravdi, C. Osmolality Effects on CHO Cell Growth, Cell Volume, Antibody Productivity and Glycosylation. Int. J. Mol. Sci. 22, 3290 (2021).

63. Chen, P. & Harcum, S. W. Effects of amino acid additions on ammonium stressed CHO cells. J. Biotechnol. 117, 277–286 (2005).

64. Schneider, M., Marison, I. W. & von Stockar, U. The importance of ammonia in mammalian cell culture. J. Biotechnol. 46, 161–185 (1996).

65. Buchsteiner, M., Quek, L.-E., Gray, P. & Nielsen, L. K. Improving culture performance and antibody production in CHO cell culture processes by reducing the Warburg effect. Biotechnol. Bioeng. 115, 2315–2327 (2018).

66. Gupta, S. K. et al. Metabolic engineering of CHO cells for the development of a robust protein production platform. PLOS ONE 12, e0181455 (2017).

67. Romanova, N. et al. Hyperosmolality in CHO culture: Effects on cellular behavior and morphology. Biotechnol. Bioeng. 118, 2348–2359 (2021).

68. Pan, X., Dalm, C., Wijffels, R. H. & Martens, D. E. Metabolic characterization of a CHO cell size increase phase in fed-batch cultures. Appl. Microbiol. Biotechnol. 101, 8101–8113 (2017).

69. Kiehl, T. R., Shen, D., Khattak, S. F., Jian Li, Z. & Sharfstein, S. T. Observations of cell size dynamics under osmotic stress. Cytometry A 79A, 560–569 (2011).

70. Jiang, Y. et al. The Prognostic Role of Cyclin D1 in Multiple Myeloma: A Systematic Review and Meta-Analysis. Technol. Cancer Res. Treat. 21, 15330338211065252 (2022).

71. Edagawa, M. et al. Role of Activating Transcription Factor 3 (ATF3) in Endoplasmic Reticulum (ER) Stress-induced Sensitization of p53-deficient Human Colon Cancer Cells to Tumor Necrosis Factor (TNF)-related Apoptosis-inducing Ligand (TRAIL)-mediated Apoptosis through Up-regulation of Death Receptor 5 (DR5) by Zerumbone and Celecoxib. J. Biol. Chem. 289, 21544–21561 (2014).

72. Zhao, J., Li, X., Guo, M., Yu, J. & Yan, C. The common stress responsive transcription factor ATF3 binds genomic sites enriched with p300 and H3K27ac for transcriptional regulation. BMC Genomics 17, 335 (2016).

73. Hetz, C. The unfolded protein response: controlling cell fate decisions under ER stress and beyond. Nat. Rev. Mol. Cell Biol. 13, 89–102 (2012).

74. Jiang, H. et al. Xbp1s-Ddit3 promotes MCT-induced pulmonary hypertension. Clin. Sci. 135, 2467–2481 (2021).

75. Bustany, S., Cahu, J., Guardiola, P. & Sola, B. Cyclin D1 sensitizes myeloma cells to endoplasmic reticulum stress-mediated apoptosis by activating the unfolded protein response pathway. BMC Cancer 15, 262 (2015).

76. Varabyou, A., Salzberg, S. L. & Pertea, M. Effects of transcriptional noise on estimates of gene and transcript expression in RNA sequencing experiments. Genome Res. 31, 301–308 (2021).

77. Li, X. & Wang, C.-Y. From bulk, single-cell to spatial RNA sequencing. Int. J. Oral Sci. 13, 1–6 (2021).

78. Gyanchandani, R. et al. Intratumor Heterogeneity Affects Gene Expression Profile Test Prognostic Risk Stratification in Early Breast Cancer. Clin. Cancer Res. 22, 5362–5369 (2016).

79. Ogata, N., Nishimura, A., Matsuda, T., Kubota, M. & Omasa, T. Single-cell transcriptome analyses reveal heterogeneity in suspension cultures and clonal markers of CHO-K1 cells. Biotechnol. Bioeng. 118, 944–951 (2021).

80. Borsi, G. et al. Single-cell RNA sequencing reveals homogeneous transcriptome patterns and low variance in a suspension CHO-K1 and an adherent HEK293FT cell line in culture conditions. J. Biotechnol. 364, 13–22 (2023).

81. Shaffer, S. M. et al. Rare cell variability and drug-induced reprogramming as a mode of cancer drug resistance. Nature 546, 431 (2017).

82. Romanova, N., Schelletter, L., Hoffrogge, R. & Noll, T. Hyperosmolality in CHO cell culture: effects on the proteome. Appl. Microbiol. Biotechnol. 106, 2569–2586 (2022).

83. Moriya, H. Quantitative nature of overexpression experiments. Mol. Biol. Cell 26, 3932–3939 (2015).

84. Karottki, K. J. la C., et al. Awakening dormant glycosyltransferases in CHO cells with CRISPRa. Biotechnol. Bioeng. 117, 593–598 (2020).

85. Jiang, Q. et al. Overexpression of GRP78 enhances survival of CHO cells in response to serum deprivation and oxidative stress. Eng. Life Sci. 17, 107–116 (2016).

86. Haiyong, H. RNA Interference to Knock Down Gene Expression. Methods Mol. Biol. Clifton NJ 1706, 293–302 (2018).

87. Xiong, K. et al. Reduced apoptosis in Chinese hamster ovary cells via optimized CRISPR interference. Biotechnol. Bioeng. 116, 1813–1819 (2019).

88. Levine, M. E. & Crimmins, E. M. A Genetic Network Associated With Stress Resistance, Longevity, and Cancer in Humans. J. Gerontol. A. Biol. Sci. Med. Sci. 71, 703–712 (2016).

89. Soo, S. K. et al. Genetic basis of enhanced stress resistance in long-lived mutants highlights key role of innate immunity in determining longevity. Aging Cell 22, e13740 (2022).

90. Kourtis, N. & Tavernarakis, N. Cellular stress response pathways and ageing: intricate molecular relationships. EMBO J. 30, 2520–2531 (2011).

91. González, C. et al. Stress-response balance drives the evolution of a network module and its host genome. Mol. Syst. Biol. 11, 827 (2015).

92. Zustiak, M. P., Jose, L., Xie, Y., Zhu, J. & Betenbaugh, M. J. Enhanced transient recombinant protein production in CHO cells through the co-transfection of the product gene with Bcl-xL. Biotechnol. J. 9, 1164–1174 (2014).

93. Orellana, C. A. et al. High-antibody-producing Chinese hamster ovary cells up-regulate intracellular protein transport and glutathione synthesis. J. Proteome Res. 14, 609–618 (2015).

94. Chevallier, V., Andersen, M. R. & Malphettes, L. Oxidative stress-alleviating strategies to improve recombinant protein production in CHO cells. Biotechnol. Bioeng. 117, 1172– 1186 (2020).

95. Fabrizio, P., Garvis, S. & Palladino, F. Histone Methylation and Memory of Environmental Stress. Cells 8, 339 (2019).

96. D’Urso, A. et al. Set1/COMPASS and Mediator are repurposed to promote epigenetic transcriptional memory. eLife 5, e16691 (2016).

97. Light, W. H. et al. A Conserved Role for Human Nup98 in Altering Chromatin Structure and Promoting Epigenetic Transcriptional Memory. PLOS Biol. 11, e1001524 (2013).

98. Pascual-Garcia, P., Little, S. C. & Capelson, M. Nup98-dependent transcriptional memory is established independently of transcription. eLife 11, e63404 (2022).

99. Gialitakis, M., Arampatzi, P., Makatounakis, T. & Papamatheakis, J. Gamma Interferon-Dependent Transcriptional Memory via Relocalization of a Gene Locus to PML Nuclear Bodies. Mol. Cell. Biol. 30, 2046–2056 (2010).

100. Rupp, O. et al. A reference genome of the Chinese hamster based on a hybrid assembly strategy. Biotechnol. Bioeng. 115, 2087–2100 (2018).

101. Singh, A. & Saint-Antoine, M. Probing transient memory of cellular states using single-cell lineages. Front. Microbiol. 13, (2023).

102. Singh, A. & Hespanha, J. P. Stochastic hybrid systems for studying biochemical processes. Philos. Trans. R. Soc. Math. Phys. Eng. Sci. 368, 4995–5011 (2010).

103. Zhu, X.-F., et al. Knockdown of heme oxygenase-1 promotes apoptosis and autophagy and enhances the cytotoxicity of doxorubicin in breast cancer cells. Oncol. Lett. 10, 2974– 2980 (2015).

104. Funes, S. C. et al. Naturally Derived Heme-Oxygenase 1 Inducers and Their Therapeutic Application to Immune-Mediated Diseases. Front. Immunol. 11, (2020).

105. Orellana, C. A. et al. RNA-Seq Highlights High Clonal Variation in Monoclonal Antibody Producing CHO Cells. Biotechnol. J. 13, 1700231 (2018).

106. Pavón, M. A. et al. Enhanced cell migration and apoptosis resistance may underlie the association between high SERPINE1 expression and poor outcome in head and neck carcinoma patients. Oncotarget 6, 29016 (2015).

107. Zhang, Q., Lei, L. & Jing, D. Knockdown of SERPINE1 reverses resistance of triple-negative breast cancer to paclitaxel via suppression of VEGFA. Oncol. Rep. 44, 1875– 1884 (2020).

108. Pavón, M. A. et al. uPA/uPAR and SERPINE1 in head and neck cancer: role in tumor resistance, metastasis, prognosis and therapy. Oncotarget 7, 57351–57366 (2016).

109. Zhou, L. et al. Association of gene polymorphisms of FV, FII, MTHFR, SERPINE1, CTLA4, IL10, and TNFalpha with pre-eclampsia in Chinese women. Inflamm. Res. 65, 717–724 (2016).

110. Ma, L. & Chung, W. K. Quantitative Analysis of Copy Number Variants Based on Real-Time LightCycler PCR. Curr. Protoc. Hum. Genet. Editor. Board Jonathan Haines Al 80, 7.21.1–7.21.8 (2014).

111. Jin, H. et al. IER3 is a crucial mediator of TAp73β-induced apoptosis in cervical cancer and confers etoposide sensitivity. Sci. Rep. 5, 8367 (2015).

112. Shahid, M. et al. Emerging Potential of Immediate Early Response Gene X-1 in Cardiovascular and Metabolic Diseases. J. Am. Heart Assoc. 7, e009261 (2018).

113. Zhou, Q. et al. Dysregulated IER3 Expression is Associated with Enhanced Apoptosis in Titin-Based Dilated Cardiomyopathy. Int. J. Mol. Sci. 18, 723 (2017).

114. Arlt, A. & Schäfer, H. Role of the immediate early response 3 (IER3) gene in cellular stress response, inflammation and tumorigenesis. Eur. J. Cell Biol. 90, 545–552 (2011).

115. Zhang, J., Wang, Y., Li, G., Yu, H. & Xie, X. Down-Regulation of Nicotinamide N-methyltransferase Induces Apoptosis in Human Breast Cancer Cells via the Mitochondria-Mediated Pathway. PLOS ONE 9, e89202 (2014).

116. Xie, X. et al. Nicotinamide N-methyltransferase enhances the capacity of tumorigenesis associated with the promotion of cell cycle progression in human colorectal cancer cells. Arch. Biochem. Biophys. 564, 52–66 (2014).

117. Somerville, T. D. D. et al. TP63-Mediated Enhancer Reprogramming Drives the Squamous Subtype of Pancreatic Ductal Adenocarcinoma. Cell Rep. 25, 1741–1755.e7 (2018).

118. Melino, G. p63 is a suppressor of tumorigenesis and metastasis interacting with mutant p53. Cell Death Differ. 18, 1487–1499 (2011).

119. Chen, K. et al. Genetic analysis of heterogeneous sub-clones in recombinant Chinese hamster ovary cells. Appl. Microbiol. Biotechnol. 101, 5785–5797 (2017).

120. Chevallier, V., Schoof, E. M., Malphettes, L., Andersen, M. R. & Workman, C. T. Characterization of glutathione proteome in CHO cells and its relationship with productivity and cholesterol synthesis. Biotechnol. Bioeng. 117, 3448–3458 (2020).

121. Ku, H.-C. & Cheng, C.-F. Master Regulator Activating Transcription Factor 3 (ATF3) in Metabolic Homeostasis and Cancer. Front. Endocrinol. 11, (2020).

